# mPD5, a peripherally restricted PICK1 inhibitor for treating chronic pain

**DOI:** 10.1101/2023.03.03.530471

**Authors:** Kathrine Louise Jensen, Nikolaj Riis Chistensen, Carolyn Marie Goddard, Sara Elgaard Jager, Ida Buur Kanneworff, Alexander Jakobsen, Gith Noes-Holt, Lucía Jiménez-Fernández, Emily G. Peck, Line Sivertsen, Raquel Comaposada Baro, Grace Anne Houser, Felix Paul Mayer, Marta Diaz-delCastillo, Marie Løth Topp, Chelsea Hopkins, Cecilie Dubgaard Thomsen, Ahmed Barakat Ibrahim Soltan, Frederik Grønbæk Tidemand, Lise Arleth, Anne-Marie Heegaard, Andreas Toft Sørensen, Kenneth Lindegaard Madsen

## Abstract

Chronic pain is a complex, debilitating, and escalating health problem worldwide, impacting one in five adults. Current treatment is compromised by dose-limiting side effects including high abuse liability, loss of ability to function socially and professionally, fatigue, drowsiness, and apathy. PICK1 has emerged as a promising target for the treatment of chronic pain conditions. Here, we develop and characterize a cell-permeable fatty acid conjugated bivalent peptide inhibitor of PICK1 and assess its effects on acute and chronic pain. The myristoylated myr-NPEG_4_-(HWLKV)_2_, (mPD5), self-assembles into core-shell micelles that provide favourable pharmacodynamic properties and relieves ongoing and evoked mechanical hypersensitivity, thermal hypersensitivity as well as anxio-depressive symptoms in mouse models of neuropathic and inflammatory pain following subcutaneous administration. No overt no side effects were associated with mPD5 administration, and it has no effect on acute nociception. Finally, neuropathic pain is relieved far into the chronic phase (18 weeks post SNI surgery) and while the effect of a single injection ceases after a few hours, repeated administration provides pain relief lasting up to 20 hours after the last injection.

## INTRODUCTION

The International Classification of Diseases 11^th^ Revision (ICD-11) defines chronic pain (MG30) as a multifactorial syndrome of pain persisting for more than three months with psychological, biological, and social factors contributing to the syndrome (1). Despite complying with recommended treatment guidelines, a large fraction of the 1.5 billion people suffering from chronic pain experience compromised quality of life with constant pain and impairment of work and social life (2, 3). First-line treatments show very low efficacy of chronic pain relief, with numbers needed to treat (NTTs) ranging from six to ten depending on the aetiology of the pain condition (3–5), while opioid-based treatments entail a significant risk of high opioid use and abuse (6). The lack of efficacy and severe dose-limiting side effects originating from their centrally modulating pain transmission of current treatments highlight an urgent need to develop more effective, non-addictive pain therapeutics.

An emerging strategy for alleviating pain is modulation of receptor trafficking by targeting specific scaffold proteins (7–10). PICK1 (Protein Interacting with C Kinase 1) is a PDZ domain containing scaffold protein enriched in the postsynaptic density of neurons, known for its role in central synaptic plasticity (11, 12) and hormone storage and release (13, 14). PICK1 interacts with a host of membrane proteins and kinases via its PDZ domain (15–17), many of which have been implicated in pain signalling (18). Notably, PICK1 regulates subcellular localization and surface expression of its interaction partners, including the GluA2 subunit of α-amino-3-hydroxy-5-methyl-4-isoxazolepropionic acid (AMPA) type glutamate receptors (AMPARs) (11) and acid-sensing ion channels (ASICs) (19). PICK1 is expressed in dorsal root ganglia (DRGs) and concentrated in lamina I and the inner lamina II of the dorsal horn (7, 10, 20, 21), areas that are all important for the transmission of painful stimuli. Based on studies in animal models using inhibitory peptides, siRNA, and knock-out mice, PICK1 has been shown to be implicated in thermal and mechanical hypersensitivity in neuropathic and inflammatory pain models, advocating PICK1 as a putative target for pharmaceutical intervention of chronic pain states (7, 8, 20–22).

Developing small molecule inhibitors of PICK1 has proven difficult (17, 23, 24) with some progress in the last decade (25–27). Our recent development of a membrane permeable, bivalent, high-affinity PICK1-inhibitor, TPD5 (8), displaying low nanomolar target affinity, represents a major leap towards a potential PICK1 targeting therapeutic. Intrathecal (i.t.) administration of TPD5 fully alleviated mechanical hypersensitivity in the spared nerve injury (SNI) model of neuropathic pain. However, TPD5 was designed to penetrate the blood brain barrier and target spinal cord plasticity, raising concerns about central side effects as is known for current treatments (6, 8, 28–30). Nonetheless, TPD5 concomitantly reduced transmission in the Lissauer’s tract, demonstrating effect on the first order DRG neurons (8). In addition, we have shown that AAV-mediated expression of similar PICK1 inhibitors, confined to DRGs, is sufficient for full pain relief (31). In recent years, the use of fatty acid modifications on peptides has emerged as a successful way to enhance both plasma stability and cell permeability of pharmaceutical peptides while also offering a benevolent toxicology profile and low CNS exposure (32–34). In the current study, we describe the development of a myristoylated lipid-conjugated peptide PICK1-inhibitor, myr-NPEG_4_-(HWLKV)_2_ (mPD5), thereby circumventing concerns raised over the safety profile for TAT-conjugated peptides for a drug intended for repeated administration in particular (35, 36) and (TAT NR2B9c, US patent 8,080,518 B2).

mPD5 demonstrated high stability, solubility, and plasma protein binding, as well as low blood brain barrier penetrance, all compatible with further drug development. Functional characterization of mPD5 following subcutaneous (s.c.) administration in mice showed robust relief of mechanical and thermal hypersensitivity in mouse models of both inflammatory and neuropathic pain in female and male mice. Moreover, mPD5 reversed anxio-depressive symptoms and ongoing pain without affecting locomotor activity or putative on target effects on learning, memory, and male fertility. Contrary to other peripherally acting pain-relieving drugs, mPD5 did not affect acute nociception. Finally, repeated administration of mPD5 gave rise to sustained relief of mechanical hypersensitivity lasting 20 hours after the last administration, advancing the molecule as a strong lead molecule in the clinic for the treatment of chronic pain conditions.

## RESULTS

### mPD5 oligomerizes into micelle structures

We have previously developed and characterized the blood brain barrier permeable bivalent PICK1 peptide inhibitor, TAT-NPEG_4_-(HWLKV)_2_ (TPD5), showing promise as a therapeutic lead for the treatment of pain (8, 22) and addiction (37). To reduce potential side effects, and to facilitate plasma stability and distribution, we introduced a C_14_ fatty acid (myristic acid, myr) instead of the cell-penetrating TAT sequence on the same scaffold, NPEG_4_-(HWLKV)_2_ (PD5) (Figure 1A), resulting in myr-NPEG_4_-(HWLKV)_2_ (mPD5) (Figure 1B). mPD5 demonstrated excellent shelf stability (Table 1) and was soluble to at least 250 mg/mL (∼130 mM) in PBS, as judged from a transparent and monophasic appearance.

**Figure 1.**
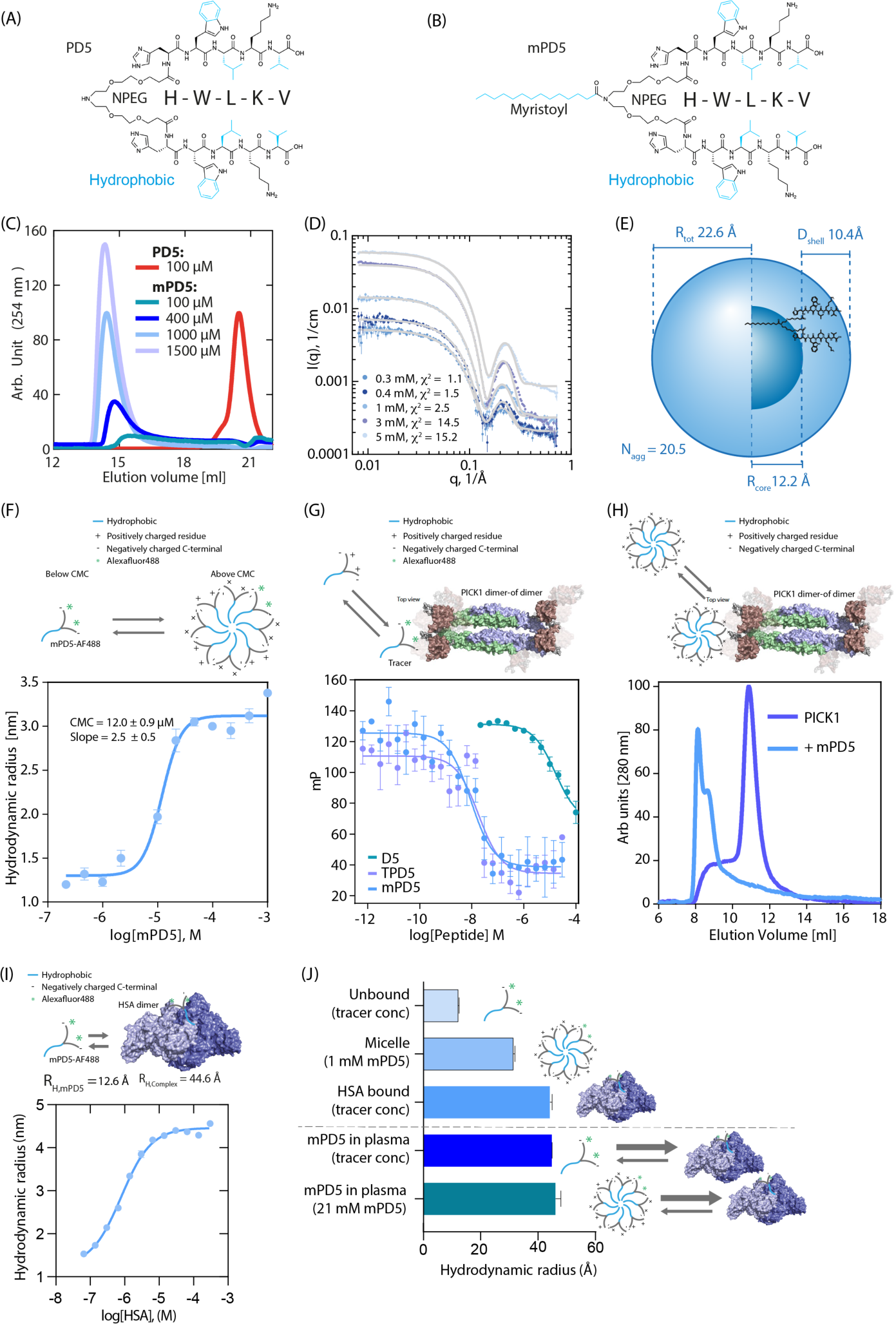
Biophysical characterization of mPD5. Structure of NPEG_4_-(HWLKV)_2_ (PD5) **(A)** and myristoyl-NPEG_4_-(HWLKV)_2_ (mPD5) **(B)** (blue: hydrophobic). **(C)** Size exclusion chromatography (SEC) with different concentrations of mPD5 demonstrating self-assembly. **(D)** Small angle X-ray scattering (SAXS) data (points) of mPD5 at different concentrations. Model fits of the core-shell model (lines) (see Supplemental Table 1 for fitting parameters). **(E)** Molecular constrained spherical core-shell model fitted to the data (Figure S1B); 20.5 molecules/micelle: total radius, R_total_ = 22.6 Å. **(F)** Flow-induced dispersion analysis (FIDA) binding isotherm of mPD5 combining optimal coatings for different concentrations (additional information in Supplemental Figure S2); CMC = 12 µM; hydrodynamic radius, *R_H_* ∼ 30 Å. **(F-I)** Experimental illustration above data. **(G)** Fluorescence polarization (FP) competition binding curves of mPD5 (blue) (K_i,app_ = 3.0 nM, SEM interval [2.3-3.8] nM, n = 6), TPD5 (purple) (K_i,app_ = 3.9 nM, SEM interval [3.5-4.4] nM, n = 3) and D5 (HWLKV) (green) (K_i,app_ = 6998 nM, SEM interval [4972-9849] nM, n = 3), using 5FAM-PD5 (5 nM vs.TPD5 and mPD5) or 5FAM-D5 (20 nM vs. D5) as tracer. Data was fitted to a competitive binding One Site fit using GraphPad prism 8.3. **(H)** SEC elution profile of PICK1 in absence (purple) or presence (blue) of mPD5, in a PICK1:mPD5 molecular ratio of 4:1. **(I)** Isotherm of mPD5-AF488 binding to human serum albumin (HSA); affinity (*K_D_*) = 787 nM; hydrodynamic radius (*R_H_*) of the complex = 4.46 nm. **(J)** Histogram showing the *R_H_*’s ±SEM of mPD5 (n = 3).

**Table 1.**
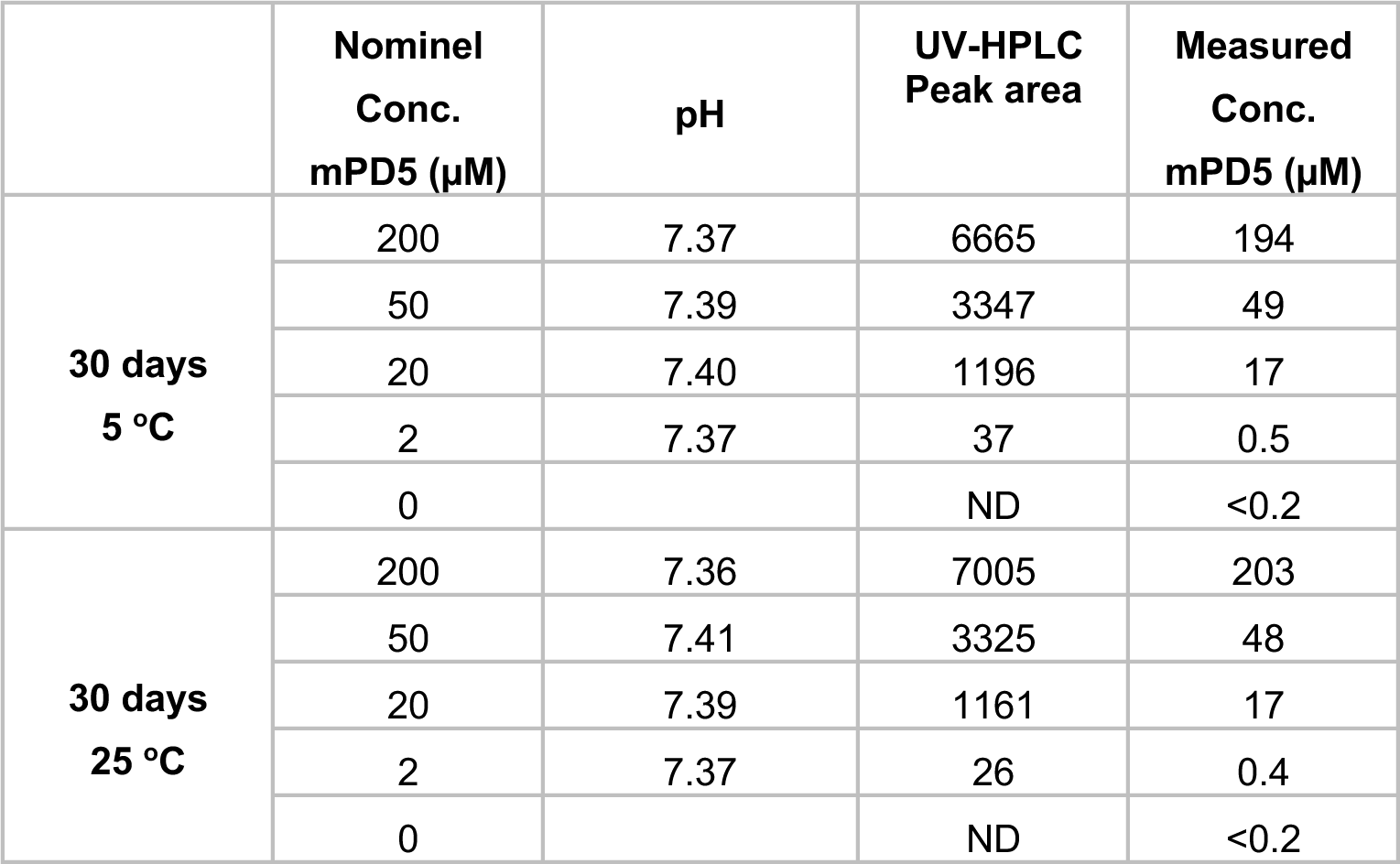
Assessment of mPD5 stability.

|  | Nominal<br>Conc.<br>mPD5 (μM) | pH | UV-HPLC<br>Peak area | Measured<br>Conc.<br>mPD5 (μM) |
| --- | --- | --- | --- | --- |
| **30 days**<br>**5 °C** | 200 | 7.37 | 6665 | 194 |
|  | 50 | 7.39 | 3347 | 49 |
|  | 20 | 7.40 | 1196 | 17 |
|  | 2 | 7.37 | 37 | 0.5 |
|  | 0 |  | ND | <0.2 |
| **30 days**<br>**25 °C** | 200 | 7.36 | 7005 | 203 |
|  | 50 | 7.41 | 3325 | 48 |
|  | 20 | 7.39 | 1161 | 17 |
|  | 2 | 7.37 | 26 | 0.4 |
|  | 0 |  | ND | <0.2 |

We hypothesized that these attractive properties might arise from an oligomeric or core-shell micellar self-assembly, with the hydrophobic lipid forming a central core and the hydrophilic PEG_4_-linked 5-mer peptide facing the aqueous solution. Using size exclusion chromatography (SEC) (Figure 1C) we found that PD5, lacking the myristic acid, eluted at ∼21 mL, whereas mPD5 eluted at ∼14.2 mL, independent of initial concentration, consistent with self-assembly of mPD5 (Figure 1C). We used Small Angle X-ray Scattering (SAXS), to investigate putative micelle structure and size. Data (Figure 1D and Supplemental Figure 1) were characteristic of core-shell micellar particles exhibiting a significant oscillation at high-*q*, as well as an extended flat Guinier region at low-*q* values, indicating small micelles without presence of larger aggregates. Further analysis of the low-*q* region showed that the forward scattering (I_0_) of mPD5 scaled linearly with concentration (Suplemental Figure S1A) suggesting concentration independent particle size with no significant interparticle interactions in the studied concentration range (0.3 - 5 mM). The pair-distance distribution, *p(r)*, showed a radius of gyration, R_g_ ∼30 Å and a D_max_ of ∼50 Å (Supplemental Figure S1C) with a shape consistent with core-shell particles (38, 39). We fitted the SAXS data using a molecularly constrained model for spherical core-shell micelles composed of an inner core of the fatty acid part of mPD5 and an outer shell of the peptide part of mPD5 (Supplemental Figure S1B and Figure 1E) (full model account in Methods). The fitting suggested that on average ∼20 mPD5 molecules make up the assembled micelle with a hydrophobic core radius of *R_core_* = 12.2 Å, a hydrophobic shell thickness of *D_shell_* = 10.4 Å and hence a total radius, *R_Total_* = 22.6 Å (Figure 1E and Supplementary Table S1). To determine the critical micelle concentration (CMC) of mPD5, we used flow-induced dispersion analysis (FIDA). FIDA suggested a CMC of 12 µM, and a hydrodynamic radius (R_H_) of ∼ 30 Å (Figure 1F and Supplemental Figure S2).

### mPD5 binds PICK1 with high affinity

Using fluorescence polarization competition binding, we determined the affinity (K_i,app_) of mPD5 for PICK1 to be 3.0 nM (SEM interval [2.3-3.8] nM), which is similar to TPD5 (K_i,app_ = 3.9 nM, SEM interval [3.5-4.4] nM) (8), demonstrating a ∼1000-fold affinity gain compared to D5 (HWLKV) alone (K_i,app_ = 6.9 µM, SEM interval [5.0-9.9] µM) and a 30-fold affinity gain compared to PD5 (K_i,app_ = 98 nM) (Figure 1G). This demonstrates that both bivalency and the lipid chain contribute to the overall binding strength. Finally, to evaluate the ability of micellar mPD5 to bind PICK1, we incubated recombinant full length PICK1 (40 µM) with mPD5 in a concentration corresponding to the CMC (10 µM) and observed a distinct shift in the peak elution volume of PICK1 from 10.9 mL in absence of mPD5 to two peaks at 8 mL and 9 mL in presence of mPD5 (Figure 1H), suggesting the ability of mPD5 to induce higher order complexes.

### mPD5 binds to human serum albumin in plasma

Fatty acids are reported to mediate drug binding to serum albumin, thereby enhancing plasma lifetime due to reduced renal clearance and metabolism (32, 33). Since mPD5 is lipidated and self assembles into micelles, we assessed if the self-assembly properties were dominant in plasma, or if the fatty acid binding to human serum albumin (HSA) could compete for mPD5 oligomerization in plasma. To this end, we first measured the binding of fluorescent mPD5 (mPD5-AF488) to HSA through FIDA (Figure 1I and Supplemental Figure S3) and found an affinity of mPD5-AF488 for HSA of 787 nM, which is 15-fold higher than the critical micellar concentration (CMC) (Figure 1F). *R_H_* further suggests that mPD5 binds to dimeric HSA (Figure 1I). To evaluate if mPD5 favours self-assembly or HSA binding in plasma, we incubated mPD5-AF488 in different concentrations of human plasma and obtained *R_H_*’s suggesting binding to HSA is favoured over micelle formation in plasma, even in a concentration of mPD5 (21 mM) that is 2000-fold above CMC (Figure 1J and Supplemental Figure S4). Taken together, our data suggest that the lipid chain drives micellar assembly of mPD5 allowing for high solubility and good stability, as well as high-affinity PICK1 binding, while once in plasma, mPD5 preferentially binds to serum albumin at the given concentrations.

### mPD5 distributes to DRGs, but not central nervous system

To assess pharmacokinetic properties of mPD5 and to guide dosing, we assessed dose dependence of the plasma exposure of mPD5 following s.c. administration using a five-fold descending dose range (50, 10, 2 μmol/kg) (Figure 2A). Plasma levels were assessed after 0.5, 1, 2, 5 and 12 hours. For all doses, we observed an initial increase in plasma concentration reaching maximal concentration after one hour (1.4 ± 0.1 mg/mL after 2 µmol/kg injection; 6.2 ± 5 mg/mL after 10 µmol/kg injection; 20.2 ± 0.6 mg/mL after 50 µmol/kg injection) followed by a linear elimination phase on the semi-log scale indicating first order kinetics. The maximal dose and area under the curve both scaled linearly with dose, and T_1/2_ showed a tendency to increase with increasing dose (0.50 ±. 0.07 hour after 2 µmol/kg injection; 0.59 ± 0.07 hour after 10 µmol/kg injection; 0.84 ± 0.03 hour after 50 µmol/kg injection). For all concentrations, the distribution volume was approximately 30 mL indicating good distribution to plasma from the site of injection.

**Figure 2.**
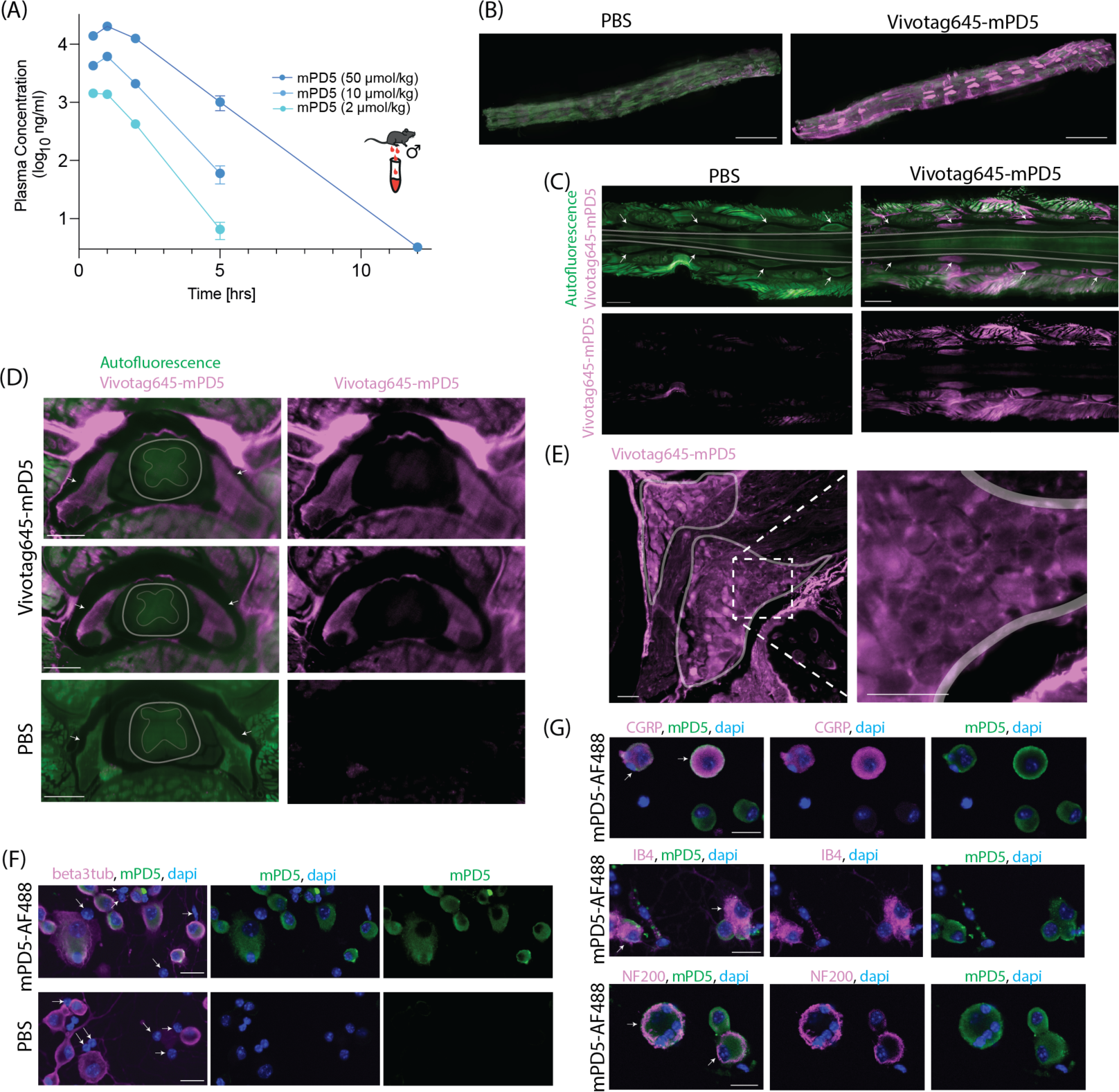
mPD5 distribution. **(A)** LC-MS/MS analysis of mPD5 plasma concentration at 0.5, 1, 2, 5 and 12 hours following subcutaneous injection in fasted male mice. Concentration peaks at one hour post injection in a dose-dependent manner (2 µmol/kg = 1.4 ±0.1 mg/mL; 10 µmol/kg = 6.2 ±5 mg/mL; 50 µmol/kg = 20.2 ±0.6 mg/mL). mPD5 is eliminated with linear kinetics. n = 3. Lower limit of quantification = 2 ng/mL. **(B)** Maximum projection of 3D imaged cleared spinal column with Vivotag645-mPD5 (magenta) and autofluorescence (green). Orientation: Caudal-rostral in the left-right direction and dorsal side facing up. Scale bar = 5000 μm. n = 3. **(C)** Optical section of 3D imaged lumbar spinal column in horizontal view with Vivotag645-mPD5 (magenta) and autofluorescence (green). Orientation: Caudal-rostral in the left-right direction and on the level of the DRGs in the dorsal-ventral direction. Arrows point to DRGs. Scale bar = 1000 μm. n = 3. **(D)** Optical section of 3D imaged spinal column in transverse view with Vivotag645-mPD5 (magenta) and autofluorescence (green). Scale bar = 500 μm Arrows point to DRGs. Grey area surrounds spinal column. n = 3. **(E)** Optical section of high-resolution light sheet imaging of one DRG. The two marked areas highlight regions with many neuronal cell bodies. The white dashed box indicates the magnified view. Scale bars = 50 μm. **(F)** Primary dorsal root ganglia culture stained against neurons with beta 3 tubulin (magenta), mPD5-488 (green) and nuclei (blue). Arrows point to non-neuronal cells with no mPD5 signal. Scale bar = 20 μm. **(G)** Primary dorsal root ganglia culture stained against neuronal subtype markers CGRP, IB4 or NF200 (magenta), mPD5-488 (green) and nuclei (blue). Arrows point to double positive cells for neuronal subtype marker and mPD5. Scale bar = 20 μm.

To determine the distribution within the nervous system, PBS or 10 µmol/kg Vivotag645-mPD5 (Figure 2B) was injected s.c. one hour before transcardial perfusion followed by dissection of brain and spinal column for whole tissue clearing and light sheet microscopy. Vivotag645-mPD5 showed distinct distribution within the spinal column (Figure 2C). Optical section providing a horizontal view of the spinal column in the plane of the DRGs revealed distribution of Vivotag645-mPD5 to striated muscle surrounding the spinal column and to DRGs of the nervous system (Figure 2D). Surprisingly, little if any signal was detected in the spinal cord suggesting exclusion by the blood brain barrier. Transverse optical section further supported this finding and highlighted high concentration of Vivotag645-mPD5 in patched structures on the dorsal part of the spinal canal (Figure 2E). Maximum projection of 3D imaged cleared whole brains revealed Vivotag645-mPD5 signal in vascular and connective tissues (Supplemental Figure S5A) but with little if any signal within the brain tissue as viewed in the sagittal plane corroborating poor blood brain barrier penetrance (Supplemental Figure S5A). In accordance, we did not detect mPD5 in CSF, spinal cord, and brain tissue by mass spectrometry following s.c. injection of mPD5 in mice (Table 2). High-resolution light sheet imaging of a single DRG suggested uptake of Vivotag645-mPD5 in somas of DRG neurons (Figure 2E). To confirm cellular uptake in DRG neurons, primary DRG cultures from adult mice were incubated with mPD5-AF488 (10 µM). Confocal imaging demonstrated cytosolic mPD5-AF488 signal surrounding the nuclei of neurons identified by beta-3 tubulin staining with no mPD5-AF488 signal in non-neuronal cells (dapi positive, beta-3 tubulin negative) (Figure 2F and Supplemental Figure S6A). Co-staining with markers of neuronal subtypes did not indicate subtype selectivity (Figure 2G and Supplemental Figure S6B).

**Table 2.**
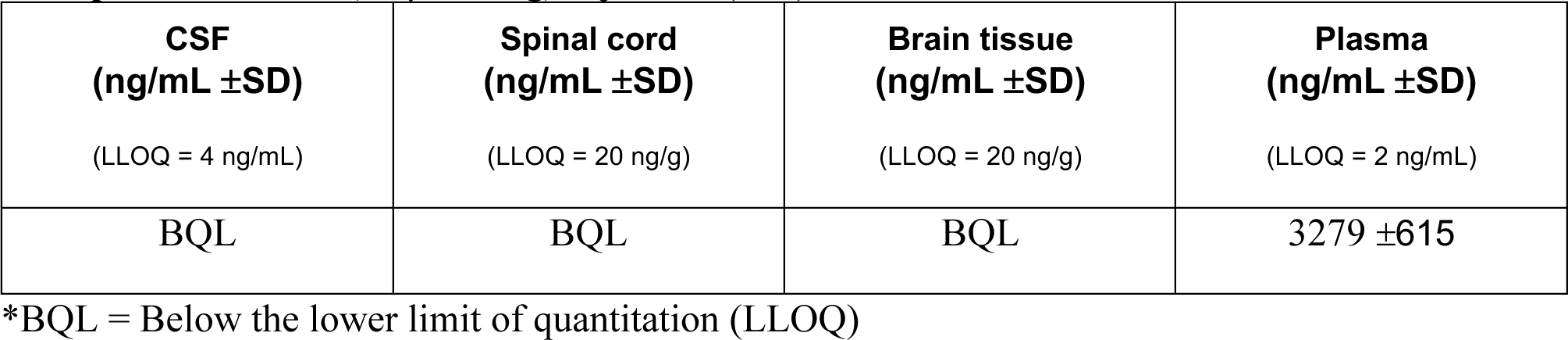
Mass spectrometry assessment of mPD5 levels in CSF, spinal cord and brain tissue one hour post s.c. mPD5 (10 µmol/kg) injection (n=3).

### Subcutaneous administration of mPD5 reduces mechanical and thermal hypersensitivity in a mouse model of inflammatory pain

We tested the effect of i.t. administered mPD5, similar to what was done with TPD5 (8), using the complete Freund’s adjuvant (CFA) model of inflammatory pain (Figure 3A). The experiment was performed as follows: on day 0, the baseline mechanical paw withdrawal threshold (PWT) was established before intraplantar injection with CFA which gives rise to a behavioural indication of allodynia reversing after 11 days, consistent with previous studies (20, 40) (Figure 3, A and B). Mice injected with saline instead of CFA (sham), showed no changes in their PWT (Figure 3B). On day two after CFA injection and following randomization, mice were injected with mPD5 (i.t., 20 µM, 7µl) resulting in a significant relief of mechanical hypersensitivity at one hour and five hours after administration, with no effect 24 hours after administration (Figure 3A). The efficacy of mPD5 by this route of administration was very similar to TPD5, confirming effects on spinal transmission, as was duration of action albeit with a slightly faster onset kinetics (8).

**Figure 3.**
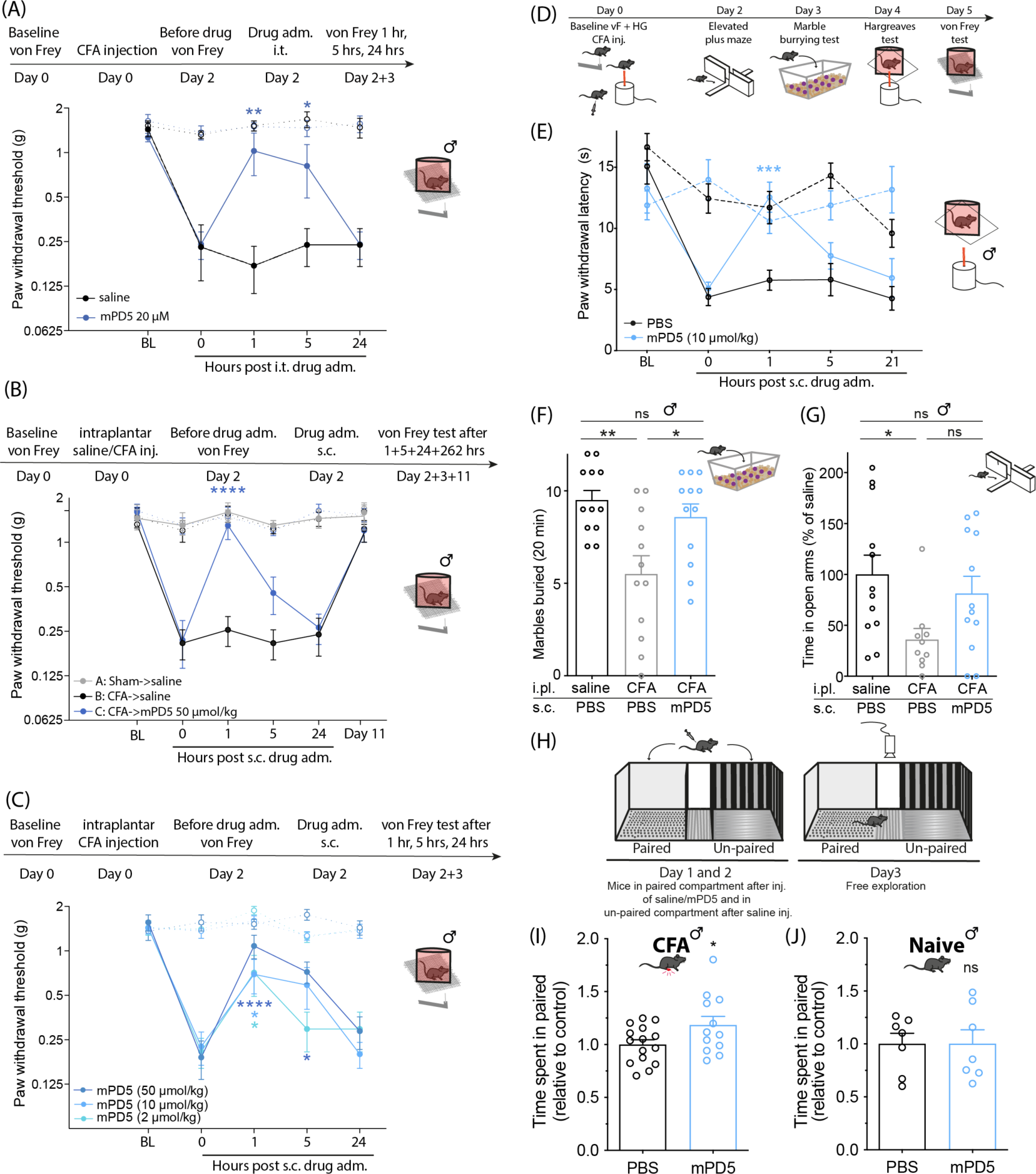
Efficacy of mPD5 in a mouse model of inflammatory pain. **(A)** Paw withdrawal threshold (PWT) before and after induction of inflammatory pain (CFA-induced) and treatment (i.t.) with mPD5 or saline. n_saline_= 5, n_mPD5_ = 6. **(B)** PWT before and after induction of inflammatory pain or sham and s.c. treatment with mPD5 or saline. n_saline->saline_ = 4, n_CFA->saline_ = 5, n_CFA->mPD5_ = 6. **(C)** PWT before and after induction of inflammatory pain and s.c. treatment with mPD5. n_50_ = 5, n_10_ = 6, n_2_ = 5. **(D)** Timeline of (E-G). **(E)** Paw withdrawal latency (PWL) before and after induction of inflammatory pain and s.c. treatment with mPD5 or PBS. n= 6. **(F)** Marbles buried after induction of inflammatory pain or sham one hour post s.c. treatment with mPD5 or PBS. n = 12. **(G)** Time spent in open arms of an elevated plus maze relative to sham, after induction of inflammatory pain or sham one hour post s.c. treatment with mPD5 or PBS. n = 12. **(H)** Schematic overview of sePP with two counter-balanced days of conditioning followed by a test day. **(I)** Time spent in the paired compartment of mice in inflammatory pain conditioned with 30 μmol/kg mPD5 or saline in the paired compartment. n_PBS_= 13, n_mPD5_ = 12. **(J)** Time spent in the paired compartment of naive mice conditioned with 30 μmol/kg mPD5 or saline in the paired compartment. n = 7. **Abbreviations;** adm. = administration, BL = baseline, CFA = Complete Freund’s Adjuvant, hrs = hours, inj. = injection, i.pl. = intraplantar, i.t. = intrathecal, s.c. = subcutaneous, sePP = single exposure place preference. Dashed line in A-C+E = contralateral paw. Statistics: A-B and E: Two-way ANOVA with Dunnetts posthoc test vs. 0 hours. F+G: One-way ANOVA with Tukey test. I+J: Unpaired t-test.

Encouraged by the biodistribution followed by s.c. administration (Figure 2), we tested the effect of mPD5 given by this route of administration in the CFA model of inflammatory pain in male mice (Figure 3, B-J). A single s.c. administered high dose (50 μmol/kg) mPD5 reversed hypersensitivity one hour after injection compared to before injection (Figure 3B). Next, we tested dose dependence of mPD5 in the CFA model using the doses tested for plasma concentrations (s.c., 2, 10 and 50 μmol/kg, 10 μl/g) (Figure 3C). All doses tested revealed significant reversal of mechanical hypersensitivity one hour post administration, and the 50 μmol/kg dose also showed significant reduction of mechanical hypersensitivity five hours post injection. In combination with the exposure data (Figure 2A), this indicates that a plasma concentration of approximately 1 mg/mL is sufficient to evoke a significant reduction in mechanical hypersensitivity. The ability of mPD5 to revert CFA induced mechanical hypersensitivity was confirmed in female mice (s.c., 10 μmol/kg) (Supplemental Figure S7A).

To test the effect of mPD5 on a different pain-related sensory modality, we assessed thermally evoked hypersensitivity in the CFA model using the Hargreaves test (Figure 3E). Following baseline testing, mice were injected with CFA and randomly assigned into treatment vs. vehicle groups. Intraplantar injection of CFA into the hind paw led to thermal hypersensitivity of the ipsilateral paw, a behavioural indication of thermal hyperalgesia, which was reversed by mPD5 (s.c., 10 μmol/kg) one hour after administration. No effect of either CFA or mPD5 was observed on the contralateral paw (Figure 3E).

### mPD5 reduces pain related behaviours in a mouse model of inflammatory pain

In the clinic, several comorbidities of chronic pain have been identified, including anxiety, depression, and fatigue (41). Mechanically and thermally evoked hypersensitivity do not assess aspects of ongoing pain, functional impairment, nor anxio-depressive symptoms associated with pain, all of which are clinically important pain-related symptoms (42). Therefore, we used the combination of a marble burying test, elevated plus maze, and single exposure place preference (sePP) to qualify the therapeutic relevance of mPD5 (Figure 3D and F-J). In the marble burying test (Figure 3F), CFA injection led to significantly decreased marble burying, presumably anxiogenic. This decrease was reversed significantly by treatment with 10 μmol/kg mPD5 to the level of the naive mice. In the elevated plus maze test (Figure 3G), CFA injection led to significantly decreased time spent in the open arms, presumably also reflecting an anxiogenic effect. Following mPD5 treatment (s.c., 10 μmol/kg) of the CFA-injected animals, the amount of time spent in the open arms was no longer significantly different from the naive mice. Finally, we used a sePP setup (43) to estimate the initial perception of the drug as a measure of relief of ongoing pain (Figure 3, H-J). The experiment was performed in a three-compartment apparatus with a striped and a grey compartment separated by a neutral zone (Figure 3H). CFA animals were injected with either PBS in both compartments or mPD5 (s.c., 30 μmol/kg) in the grey (paired) compartment and PBS in the striped (unpaired) compartment. Interestingly, the CFA animals treated with mPD5 spent significantly more time in the paired compartment compared to the PBS animals, indicating a positive effect of mPD5 on spontaneous/ongoing pain (Figure 3I). Due to the high abuse liability of the current chronic pain treatments (44), we repeated the experiment on naive mice to assess putative intrinsic rewarding properties of mPD5 (Figure 3J). The initial sensitivity to the rewarding properties of drugs is believed to be an important endophenotype in relation to the vulnerability to addiction and that the initial sensitivity to the rewarding properties of a specific drug is a relevant indicator of addictive properties of said drug (43, 45). Importantly, such control animals did not show any preference for mPD5 compared to PBS (Figure 3J).

### mPD5 reduces mechanical hypersensitivity in mice following spared nerve injury and streptozocin injection, but not in cancer induced bone pain

We next evaluated the pain-relieving effects of mPD5 in different neuropathic pain models. First, we investigated the effect of mPD5 in the SNI model in male mice (Figure 4A). On day 0, the baseline mechanical PWT was established, followed by SNI surgery. SNI surgery led to a significant decrease in PWT on day 7 versus baseline, a behavioural indication of mechanical allodynia. We found that mPD5 (s.c., 10 μmol/kg) significantly reduced mechanical hypersensitivity up to three hours, whereas a lower dose (s.c., 2 μmol/kg) showed no significant effect, indicating slightly lower potency in the SNI model as compared to the CFA model. Next, we assessed the treatment efficacy of mPD5 to relieve diabetic neuropathy using the streptozocin (STZ) model of type-1 diabetes in male mice (Figure 4B). On day 0, the baseline mechanical PWT was established, followed by injection of STZ. On day 7 after STZ injection, all mice presented a drastic increase in glycemia (from 197.4 +/- 4.4 to 533.5 +/- 10.4 mg/dL) validating the diabetic state of the mice. STZ injection led to a significant decrease in PWT on day 13 versus baseline, a behavioural indication of mechanical allodynia (Figure 4B). Since the STZ model affects both paws equally, we used the established pain-relieving drug pregabalin as a positive control in the experiment instead of the contralateral paw. mPD5 (s.c., 2 and 10 μmol/kg) resulted in a significant relief of mechanical hypersensitivity the first hour post injection, and this effect was extended for another hour at the highest dose. The positive control (pregabalin) resulted in significant pain relief up to four hours after injection, whereas the vehicle (PBS) had no effect.

**Figure 4.**
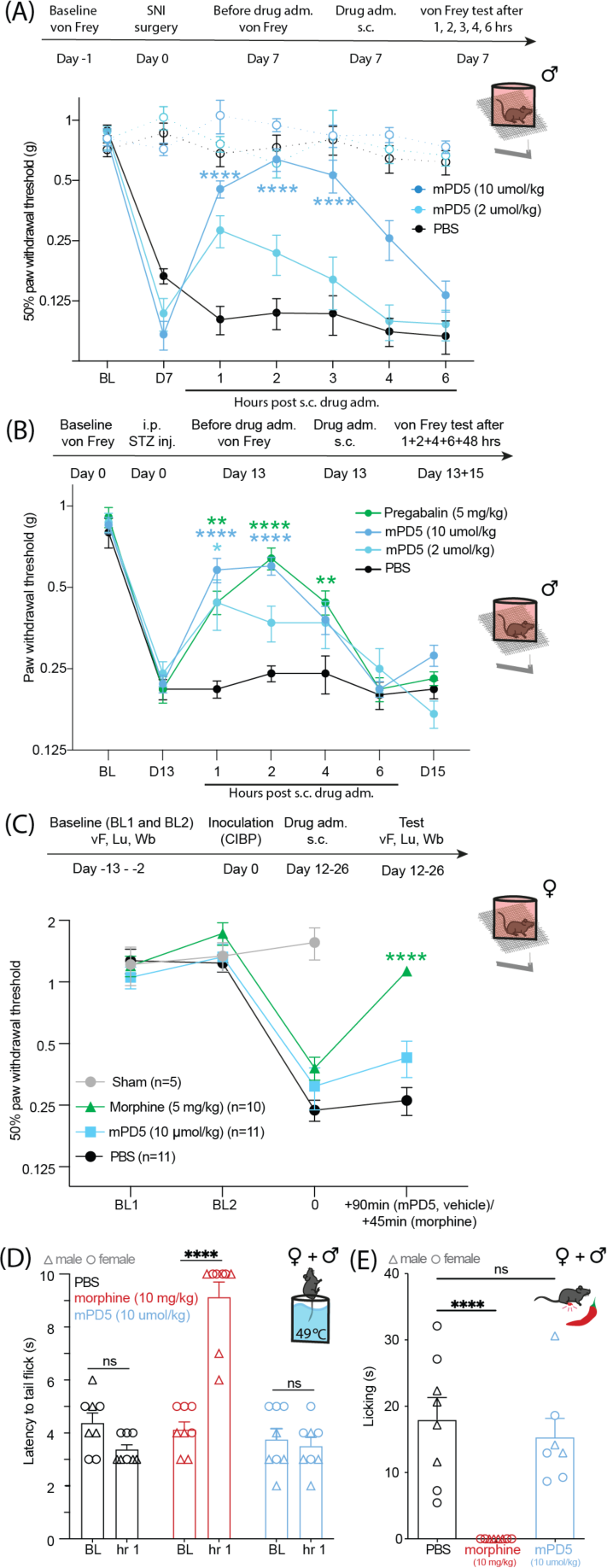
Efficacy of mPD5 in neuropathic pain, diabetic neuropathy, bone cancer induced pain and acute nociception. **(A)** Paw withdrawal threshold (PWT) before and after induction of neuropathic pain (SNI surgery) and subcutaneous (s.c.) treatment with mPD5 or PBS 7 days after surgery. n = 6 in each group. Dashed line = contralateral paw. **(B)** PWT before and after induction of diabetic neuropathy (STZ injection) and subcutaneous (s.c.) treatment with mPD5, Pregabalin or PBS 13 days after STZ injection. n_pregabalin_ = 10, n_mPD5_ = 10 and n_PBS_ = 9. **(C)** PWT before and after induction of cancer induced bone pain (sham or NCTC 2472 cell inoculation) and subcutaneous (s.c.) treatment with mPD5, morphine or PBS. n_morphine_ = 10, n_mPD5_ = 11 and n_PBS_ = 11. **(D+E)** Efficacy of mPD5 on acute pain in female (o) and male (Δ) mice. **(D)** Effect of PBS, morphine (10 mg/kg) and mPD5 (10 μmol/kg) on tail-flick time in water of 49 ±0.5 °C. **(E)** Effect of PBS, morphine (10 mg/kg) and mPD5 (10 μmol/kg) on capsaicin-induced licking time. Abbreviations; adm. = administration, BL = baseline, CIBP = cancer induced bone pain, D = day, hrs = hours, inj. = injection, s.c. = subcutaneous, stz = streptozocin, SNI = spared nerve injury. Statistics: A-C: Two-way ANOVA with Dunnetts posthoc test vs. 0 hours. D: Two-way ANOVA with Sidak posthoc test of BL vs one hour. E: One-way ANOVA with Dunnetts posthoc test of PBS vs drug.

Finally, we assessed the effect of mPD5 in a cancer induced bone pain (CIBP) model (Figure 4C and Supplemental Figure S7B-C). Mice were inoculated with a sarcoma cell line (NCTC 2472) in the femur marrow cavity. Pain-like behaviour was assessed every second day until meeting the criteria of a limb use score of two or below in combination with a weight bearing ratio of 0.35 or below, where they received treatment with mPD5 (s.c. 10 μmol/kg) or vehicle. Mice met the criteria at 12-26 days after inoculation. Since symptoms of pain are notoriously challenging to relieve in CIBP models, we used morphine as a positive control. Indeed, morphine (s.c., 5 mg/kg) gave rise to a partial but significant increase in PWT, limb use score, and weight bearing ratio in the animals, whereas mPD5 and vehicle had no effect on either.

### mPD5 has no effect on nociceptive responses in naive mice

We next assessed the effect of mPD5 in two models of acute pain in both sexes. In the hot water tail immersion test, morphine (s.c., 10 mg/kg) significantly reduced the latency to tail-flick compared to baseline, whereas PBS and mPD5 (s.c., 10 μmol/kg) had no effect on tail-flick latency (Figure 4D). Similarly, morphine (s.c., 10 mg/kg) significantly reduced the licking time of mice following intraplantar (i.pl.) injection of capsaicin, whereas PBS and mPD5 (s.c., 10 μmol/kg) had no effect on time spent licking (Figure 4E).

### mPD5 reduces pain related behaviours in a mouse model of neuropathic pain

To explore further the effect of mPD5 on neuropathic pain, we returned to the combination of von Frey, marble burying test, and elevated plus maze; this time in female mice using the SNI model (Figure 5A). Using von Frey, we found that mPD5 significantly reduced mechanical hypersensitivity one hour post injection at all tested doses (s.c., 2, 10, 50 μmol/kg) with no effect of PBS (Figure 5B). In the marble burying test (Figure 5C), SNI surgery led to significantly decreased marble burying, and this decrease was reversed significantly by mPD5 treatment (s.c., 10 μmol/kg) to the level of the naive mice. In the elevated plus maze, the SNI did not affect the time spent in open arms compared to naive mice (Supplemental Figure S7D). We tested the effect of mPD5 in the sePP for the SNI model (Figure 5D) using the same setup as for the CFA model (Figure 3H). However, mPD5 (s.c., 30 μmol/kg) did not change the time spent in the paired chamber compared to PBS in the SNI model (Figure 5D). As an alternative measure of the ongoing pain perception, we recorded the ultrasonic vocalizations. SNI surgery led to significantly more vocalizations at 37 kHz, and this increase was fully reversed by mPD5 treatment (s.c., 10 μmol/kg) to the level of the naive mice (Figure 5E).

**Figure 5.**
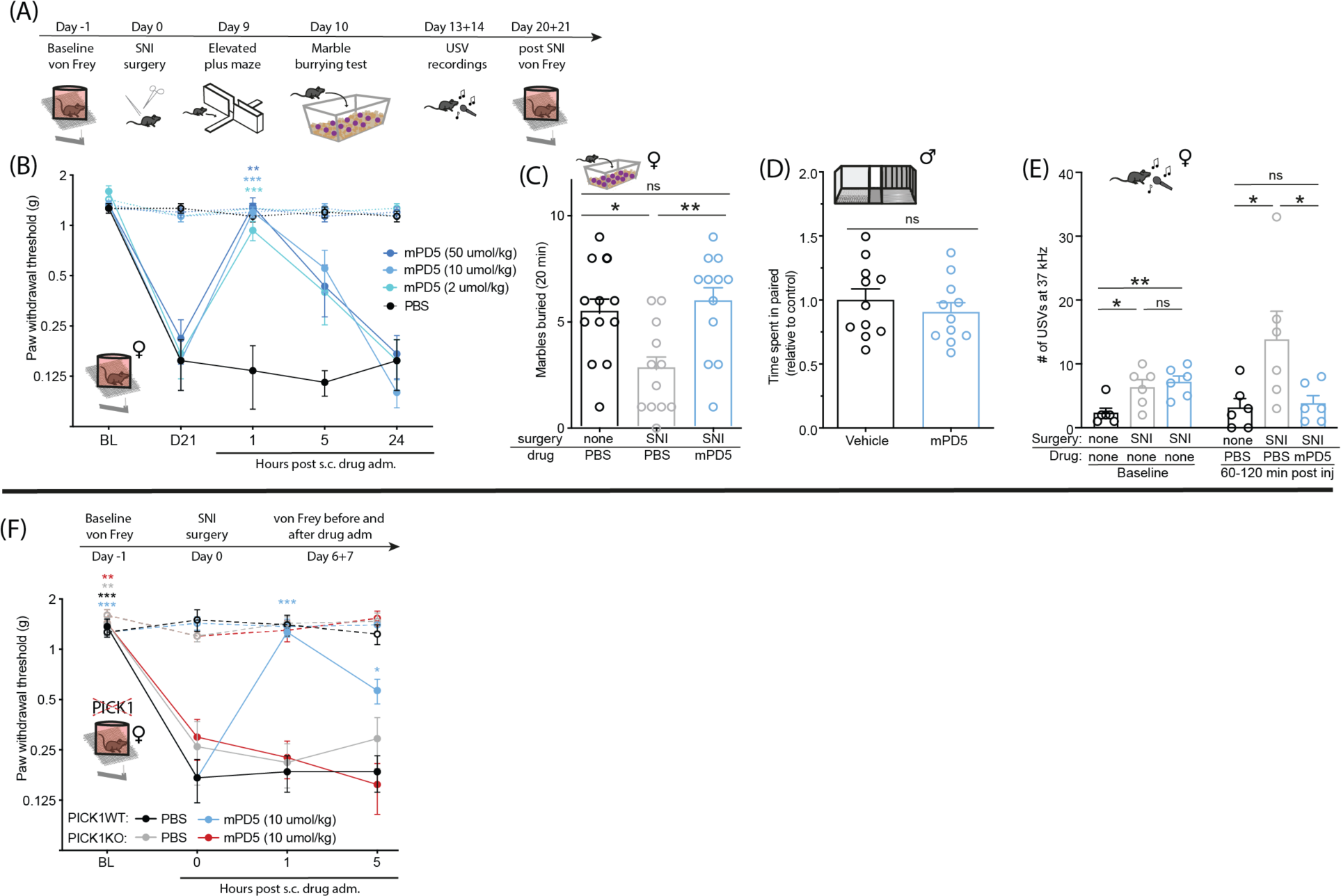
Efficacy of mPD5 in a model of neuropathic pain in wildtype and PICK1 KO mice. **(A)** Timeline for B-C and E. **(B)** Paw withdrawal threshold (PWT) before and after induction of neuropathic pain (SNI surgery) and subcutaneous (s.c.) treatment with mPD5 or PBS. n = 6. **(C)** Marbles buried after induction of neuropathic pain or naive one hour post s.c. treatment with mPD5 or PBS. n = 12 in each group. **(D)** Time spent in the paired compartment of SNI mice conditioned with 30 μmol/kg mPD5 or saline in the paired compartment. n = 11. **(E)** Recordings of Ultrasonic vocalizations of SNI and naive mice at 37 kHz made for 60 minutes at baseline and 60-120 min post subcutaneous (s.c.) treatment with mPD5 or PBS. n = 6. **(F)** PWT before and after induction of neuropathic pain and subcutaneous (s.c.) treatment with mPD5 or PBS in PICK1 wildtype and knockout mice. Cross-sectional study ending up with n = 6 in each group. Abbreviations; adm. = administration, BL = baseline, D = day, inj. = injection, s.c. = subcutaneous, SNI = spared nerve injury, USV = ultrasonic vocalizations. Statistics: B+F: Two-way ANOVA with Dunnetts posthoc test vs. 0 hours. C: One-way ANOVA with Tukey multiple comparison. D: Unpaired t-test. E: One-way ANOVA of SNI-PBS vs the other two groups at baseline and after treatment. Dashed line = contralateral paw.

### mPD5 does not revert hypersensitivity following SNI surgery in mice lacking PICK1

It has previously been shown that the hypersensitivity of PICK1 KO mice is blunted in the L5 spinal nerve ligation (SNL) model (21), in which transection is made close to the DRG of the L5 only (46). In contrast, we found that female PICK1 KO mice displayed mechanically evoked hypersensitivity indistinguishable from wildtype littermates following SNI surgery (Figure 5F), in which the common peroneal and tibial nerves are cut distal to the DRGs of the L3-L5 (46). This indicates that the plasticity induced by the two models (47) may differentially rely on PICK1. Nevertheless, treatment with mPD5 had no effect on mechanical hypersensitivity in PICK1 KO mice following SNI surgery, whereas it was significantly reduced in WT littermates, consistent with PICK1 being the target of mPD5 (Figure 5F).

### mPD5 does not affect general locomotion, fertility, or learning and memory

Drug treatments of chronic pain are compromised by dose-limiting side effects (48, 49). To assess potential generalized side effects in terms of motor function, sedation, and hyperactivity, mice of both sexes were injected with PBS or mPD5 (s.c.,10 µmol/kg) and placed in an open field for 150 min (Figure 6, A and B). A tendency towards lower locomotion was observed initially (although not significant at any individual time bin), and the overall locomotion was the same between groups. Mice injected with higher doses (s.c., 30 or 50 µmol/kg) of mPD5 also did not show any significant effect on locomotor activity (Supplemental Figure S8A).

**Figure 6.**
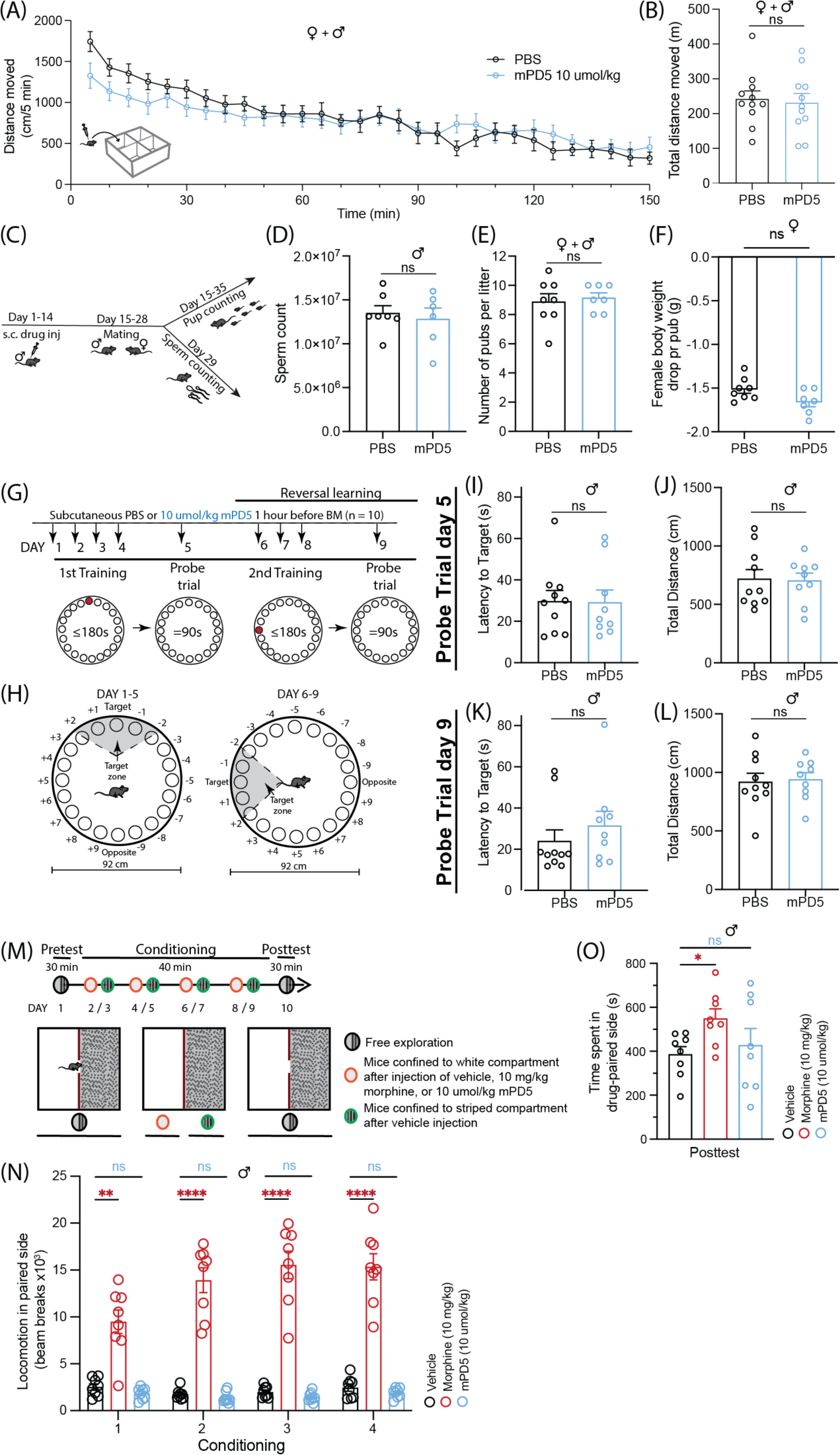
Effect of mPD5 on naive animals. **(A+B)** Locomotor response of mice injected s.c. with PBS or 10 μmol/kg mPD5 and left in open field boxes (white, 40 x 40 x 80 cm) for 150 min. Data is depicted in bins of 5 min **(A)** and as total locomotion **(B)**. (n = 11 (5 female, 6 male)). **(C)** Timeline of fertility study shown in **(D-F)**. **(D)** Sperm count (n_PBS_ = 7, n_mPD5_ = 6). **(E)** Number of pups per litter (n_PBS_ = 8, n_mPD5_ = 7). **(F)** Female body weight difference before and after labour divided by number of pups (n_PBS_ = 8, n_mPD5_ = 7). **(G-L)** Effect of mPD5 on reversal learning. **(G)** Schematic overview of the long-term retention experiment (n_PBS_ = 10, n_mPD5_ = 9). **(H)** Schematic illustration of the Barnes maze used. **(I+J)** Probe test on day five following initial four days of training **(I)** Latency to reach target hole. **(J)** Total distance moved. **(K+L)** Probe test on day nine following three days of reversal learning. **(K)** Latency to reach target hole. **(L)** Total distance moved. **(M-O)** Conditioned place preference (CPP). **(M)** Schematic overview of CPP (n=8). **(N)** Total locomotion of the three groups in the drug-paired compartment for the four days of conditioning in the white side of the CPP apparatus. **(O)** Time spent in the drug-paired compartment following conditioning. Abbreviations: BM: barnes maze, inj: injection, s.c.: subcutaneous. Statistics: A: Two-way ANOVA with Sidak multiple comparison. B+O: One-way ANOVA. I-L: Mann-Whitney t-test. N: Two-way ANOVA with Dunnets posthoc test.

Next, to address putative on-target side effects, we tested the effect of repeated administration of mPD5 (s.c., 10 µmol/kg, once daily for 14 days) vs. vehicle on male fertility (Figure 6, C-F and Supplemental Figure S8, B and C), since complete loss of PICK1 is known to cause male infertility (50). The sperm count of the males was the same between groups (Figure 6D) as was the number of pups per litter (Figure 6E and Supplemental Figure S8B). Although the weight gain was significantly higher for the females mated with mPD5-treated male mice (indicating heavier litter) (Supplemental Figure S8C), the body weight drop per pub following birth was the same for both groups (Figure 6F). In the CNS PICK1 has been strongly implicated in synaptic plasticity underlying learning and memory, as well as addictive processes (12, 18). Consequently, despite the apparent exclusion of the peptide from the CNS (Figure 2), we tested the effect of mPD5 on Barnes’ maze performance (Figure 6, G-L) as well as conditioned place preference (CPP) (Figure 6, M-O). For the Barnes’ maze experiment, mice were injected with PBS or mPD5 (s.c., 10 µmol/kg) 60 min before placement on the maze. Following four days of training (and four injections in total), the two groups showed no difference in latency to reach target (Figure 6I), or total distance moved (Figure 6J) indicating no effect on learning or recall. Likewise, no difference was observed in latency to reach the target (Figure 6K), or total distance moved (Figure 6L) following reversal learning. The overall distribution of nose pokes between the different holes also did not differ between groups in either test (Supplemental Figure S8, D and E).

### mPD5 does not show abuse liability

Finally, current chronic pain treatments, and opioids in particular, show high abuse liability (6). To compare the abuse liability of mPD5 to morphine, we performed a CPP in naive animals (Figure 6M). Morphine significantly increased locomotion during conditioning compared to PBS, whereas mPD5 did not affect locomotion in either direction (Figure 6N). As anticipated, the group treated with morphine spent significantly more time in the drug-paired compartment during the posttest compared to the PBS group, while the mPD5 group did not (Figure 6O).

### mPD5 provides sustained relief of mechanical hypersensitivity in the chronic phase of neuropathic pain

To test for putative development of tolerance due to down-regulation of the target or other adaptive mechanisms, we carried out a full dose dependence of mPD5 in the SNI model (three weeks after surgery) in mice on two consecutive days (s.c., 2, 10, 50 µmol/kg mPD5) (Figure 7A). We observed dose-dependent relief of the mechanical hypersensitivity for all concentrations tested giving rise to significant effects at one hour, while only the 50 µmol/kg showed a significant effect at five hours. Such dose-dependency was also evident the next day. Here, only the two highest concentrations showed significant pain relief, but now at both one and five hours after administration. Taken together this experiment does not indicate immediate development of tolerance.

**Figure 7.**
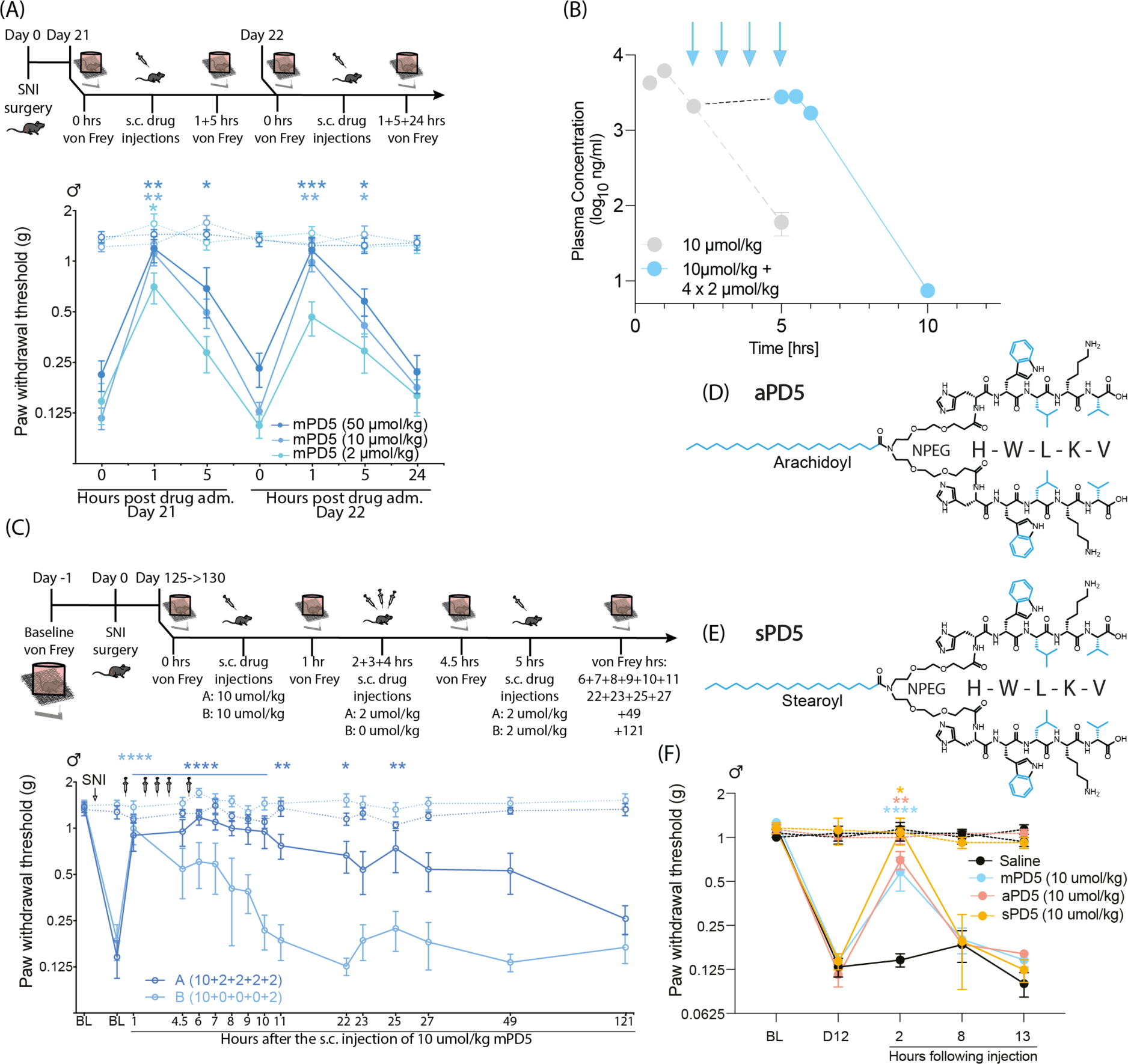
Efficacy of mPD5 on late-stage neuropathic pain following sustained dosing. **(A)** Paw withdrawal threshold (PWT) before and after induction of neuropathic pain (SNI surgery) and repeated subcutaneous (s.c.) treatment with mPD5. n = 8 in each group. **(B)** Plasma concentration of mPD5 (blue) determined by LC-MS/MS in mice injected s.c. with 10 μmol/kg mPD5 followed by 2+2+2+2 μmol/kg mPD5 once an hour. mPD5 was eliminated with linear kinetics similar to the single administration (Gray, dashed line, identical to Figure 4B), shown again for comparison. **(C)** PWT before and after induction of SNI and sustained subcutaneous (s.c.) treatment with 10 + 2 + 2+ 2 + 2 μmol/kg or 10 + 0 + 0 + 0 + 2 μmol/kg (one hour between injections) mPD5. n=8. **(D)** Structure of arachidoyl-NPEG_4_-(HWLKV)_2_ (peptide conjugated with C20 (arachidic acid)) (aPD5) and **(E)** Stearoyl-NPEG_4_-(HWLKV)_2_ (peptide conjugated with C18 (stearic acid)) (sPD5) **(F)** PWT before and after SNI surgery and s.c. treatment with saline, aPD5, sPD5 or mPD5. n = 6. Dashed line = contralateral paw. Abbreviations; adm. = administration, BL = baseline, D = day, hrs = hours, inj. = injection, s.c. = subcutaneous, PWT = paw withdrawal threshold, SNI = spared nerve injury. A+C+F: Dashed line = contralateral paw. Statistics: A+C+F: Two-way ANOVA with Dunnets posthoc test.

With this in mind, we sought to obtain sustained relief of mechanical hypersensitivity by consecutive administrations of mPD5. To estimate dose and dosing interval for maintaining a steady state plasma concentration, we employed van Rossum’s equation using parameters obtained from the single administration (D = 10 µmol/kg, Vd ∼ 30 ml, T_1/2_ = 0.6 hr). This predicted that a steady state plasma concentration of ∼2 mg/mL, following a 10 µmol/kg bolus injection, could be maintained by a dosing of 2 µmol/kg administered once an hour. We experimentally verified this prediction by assessing the plasma exposure following a single s.c. injection of 10 µmol/kg mPD5 followed by four consecutive administrations of 2 µmol/kg in one-hour intervals (Figure 7B). Blood samples were taken at 5 min, 30 min, 1 hour, 5 hours, 12 hours, 24 hours and 48 hours after the last administration. Notably, the level immediately after the last injection was 2.8 ± 0.5 mg/mL, in good agreement with model prediction. The subsequent kinetics were comparable to the single administration with a T_1/2_ of 0.52 ± 0.02 hours.

Next, we assessed how a repeated administration paradigm giving rise to a steady state plasma level of the drug would affect mechanical hypersensitivity in tbe SNI model of chronic neuropathic pain. To this end, we used mice in a very late stage of the SNI model (18 weeks post-surgery), where mice were injected with either 10 + 2 + 2 + 2 + 2 µmol/kg mPD5 (Group A) or 10 + 0 + 0 + 0 + 2 µmol/kg mPD5 as reference (Group B) (one hour between injections) (Figure 7C). Such consecutive administration of mPD5 not only showed sustained relief of mechanical hypersensitivity but surprisingly also extended the duration of effect from one hour to 25 hours (20 hours following last injection). Finally, we asked if such extended duration of action could be obtained by using peptides where the myristic acid was exchanged by longer acyl chains to increase their plasma lifetimes (51). For this, we tested peptides conjugated with C18 (stearic acid) and C20 (arachidic acid) fatty acids to produce sPD5 and aPD5, respectively (Figure 7, D-F). Single s.c. administration of sPD5 and aPD5 both caused profound itching in the mice, but nonetheless relieved mechanical hypersensitivity in the SNI model similar to mPD5 two hours after injection. Similar to mPD5, however, there was no effect on mechanical hypersensitivity eight hours after injection, suggesting that the extended effect could not be achieved by increasing the length of the fatty acyl chain only.

## DISCUSSION

In this study, we have shown that mPD5 relieved pain in three different pain models with fundamentally different aetiologies: the CFA model of acute inflammatory pain as well as the SNI and STZ models of chronic neuropathic pain. We also report that mPD5 showed no effect in a cancer induced bone pain model, where pain is known to be very challenging to treat (52). Importantly, a number of changes occur in the dorsal root ganglia, as well as the dorsal horn following induction of both inflammatory and neuropathic pain models (53). In murine models of inflammatory, neuropathic, and cancer pain it has been shown that each model generated a unique set of neurochemical changes in the spinal cord and sensory neurons (54). In C3H/Hej mice, changes in substance P and calcitonin gene-related peptide is observed in models of inflammatory (CFA) and neuropathic (sciatic nerve transection or L5 spinal nerve ligation) pain models, with no changes in levels of substance P and calcitonin gene-related peptide observed in the model of cancer pain (injection of osteolytic sarcoma cells into the femur), suggesting that cancer induces a unique persistent pain state (54). The full efficacy in neuropathic pain combined with complete lack of efficacy in cancer induced bone pain suggests that these models modify nociceptive transmission differently based on their dependence on PICK1.

We recently published on a TAT-conjugated, bivalent, high-affinity PICK1 inhibitor (TPD5) displaying robust efficacy in the SNI model of neuropathic pain and CFA model of inflammatory pain following i.t. administration in mice (8, 22). Efficacy of TPD5, however, was relatively low following systemic administration (i.p. and s.c.) and increased dosing caused significant discomfort in the mice. Symptoms included itching, respiratory abnormality, and immobility, which has been reported also for another TAT-conjugated cell-permeable peptide (TAT NR2B9c, US patent 8,080,518 B2). Of additional concern, the TAT sequence itself has been shown to alter the expression of specific genes (both induction and repression) in HeLa cells (35). In the current paper, to circumvent potential safety issues with the TAT cell-penetrating peptide, we developed mPD5, which was obtained by substitution of TAT with a C_14_ fatty acid (myristic acid). Myristoylation of simple peptides to modulate synaptic transmission and plasticity has been used previously and render them cell-permeable (7, 55, 56). In drug development, lipidation of peptides has been employed to enhance plasma stability due to the interaction between lipids and serum albumin (32, 33, 57, 58). Studies suggested that lipidation allowed the formation of higher order structures (59–61), which increased the solubility and resilience to degradation. Presumably, mPD5 benefits from all these consequences of the myristoylation, and the compound relies on well-known chemical principles and well-tested building blocks that are considered safe in humans (33, 62). In our case, the change from TPD5 to mPD5 turned out advantageous since it allowed for efficacious s.c. administration. Subcutaneous administration of peptidic drugs is more advantageous than intravenous or i.t. administration in terms of improving patient compliance, e.g. due to the suitability for self-administration (63). Subcutaneous injection is further valued due to the avoidance of hepatic and gastrointestinal degradation. Clinically, subcutaneous injection is the most common route of administration for peptides and is used extensively for both continuous and low-dose drug treatment (63–65).

It has been hypothesized that the poor translational value of preclinical data of rodents to humans, is because assessment of hypersensitivity using von Frey and Hargreaves relies on withdrawal reflexes and thus shall not stand alone (66). Our data showed that animals with inflammatory pain preferred the mPD5-paired compartment over vehicle treatment in the sePP test, indicating that the effect of mPD5 is not merely reflex inhibition. Moreover, mPD5 reduced anxio-depressive behaviours associated with pain in both inflammatory and neuropathic models, as well as reduced ultrasonic vocalizations of mice in neuropathic pain. This suggests that the mechanism of mPD5 indeed taps into the complex pattern of symptoms that are equally of relevance in chronic pain patients.

Many drugs used in the treatment of chronic pain show highly problematic central side effects including sedation, confusion, and memory problems (67, 68). Unlike opioids (67) and gabapentinoids (68), mPD5 did not significantly affect novelty induced exploration, general locomotor activity, or memory and learning. For opioids in particular there is a high risk of substance abuse (69) but abuse liability of gabapentinoids is also gaining attention (44). Indeed, morphine conferred conditioned place preference in our assessment, while the preference for the mPD5-paired compartment was not different from vehicle both following single and multiple exposures to mPD5. Since mPD5 did not reach the CNS, these experiments collectively argue that the potential abuse liability of mPD5 is low. Despite the lack of effect on the mean preference change in both experiments (Figure 3J and Figure 6O), it does seem from both experiments that the mPD5 group potentially splits into two groups. Together with the tendency to reduced locomotion (non-significant) in the exploratory phase of the open field test and our cFOS data in obese mice (70) this might warrant further studies of (indirect) effects of mPD5 on the dopamine system. Finally, contrary to other peripherally acting drugs, such as lidocaine, mPD5 relieved maladaptive pain specifically while retaining acute nociceptive and mechanical sensation. Relieving chronic pain, without limiting the sensitivity to potential harmful stimuli of everyday life (unlike i.e., morphine), would be a great benefit for patients.

In conclusion, we have shown that mPD5 relieved ongoing and evoked hypersensitivity in multiple mouse models of pain in female and male mice with cross-laboratory validation for the SNI model. mPD5 displayed favourable pharmacokinetic properties (easily soluble and highly stable). It alleviated evoked pain (thermal and mechanic) following different routes of administration (i.t. and s.c.), in inflammatory (CFA) and neuropathic pain models (SNI and STZ) and was efficacious on transient and chronic pain. Notably, and important for the translational potential of mPD5, it also reduced anxio-depressive behaviour (marble burying test and elevated plus maze) and induced place preference for the treatment-paired compartment in the inflammatory pain model.

Finally, the side effect profile of mPD5 differed substantially from the current standard of care for chronic pain conditions, including both centrally and peripherally acting drugs. Taken together, these features advocate that mPD5 represents a compelling drug candidate for further preclinical testing before clinical trials and treatment of chronic pain.

## METHODS

See supplementary information.

### Study approval

Experiments involving animals were performed in accordance with guidelines of the Danish Animal Experimentation Inspectorate (permission number 2016-15-0201-00976, 2021-15-0201-01036, 2020-15-020100439, 2022-15-0201-01216) in a fully AAALAC-accredited facility under the supervision of local animal welfare committee. In all animal experiments, the experimenter was blinded to mice treatment, except for groups treated with 10 mg/kg morphine, since the “morphine tail” gives it away.

## Data availability

Values for all data points in graphs are reported in the Supporting Data Values file.

## Author contributions

KLJ, CMG, GNH, and LS were involved in SNI experiments. KLJ performed CFA von Frey experiments and analysis on behavioural data. KLJ and LJF performed CFA anxio-depressive experiments. KLJ performed sePP and CPP experiments. KLJ performed USV experiment and EGP analysed the data. CMG performed Barnes’ maze and open field experiments. CMG and KLJ performed capsaicin and tail immersion experiments. IBK, CH, CDT, ABIS, and MDC conducted the CIBP experiment and analysed the data with KLJ. NRC and GNH performed and analysed biochemical and biophysical experiments with assistance on SAXS beamtime from FGT and model based SAXS data analysis performed by LA. RCB and LS conducted experiments. MLT created the illustrations of mPD5 and PD5. SEJ performed whole tissue clearing and immunocytochemical experiments followed by interpretation and visualization. GAH assisted with whole tissue clearing experiments. AJ performed fertility experiments with help from FPAM. KLM, ATS, and AMH supervised the research. KLJ, NRC, KLM, and ATS conceptualized the study, designed the research, and interpreted data. NRC and KLM invented the dimeric peptides. KLJ and KLM wrote the manuscript with contributions from NRC. All authors reviewed and critically evaluated the manuscript.

## Acknowledgements

We thank Morgan Thomsen and Simone Tonetto for lending us their recording equipment and their input on how to analyse ultrasonic vocalizations is greatly appreciated. We thank Nabeela Khadim for excellent technical assistance and acknowledge the Core Facility for Integrated Microscopy, Faculty of Health and Medical Sciences, University of Copenhagen. We also thank Fida Biosystems ApS, Denmark for assistance and for allowing us to conduct our experiments at their facilities and we gratefully acknowledge SAXS beam time at the P12 beamline at the PETRAIII at DESY, Germany, along with help from the beamline scientists. Lastly, we thank Zyneyro/Combigene for providing peptides. This work was supported by a Novo Nordisk Foundation Pre-seed grant (KLM, ATS), Lundbeck Foundation grants R344-2020-1063 (KLM), R322-2019-1816 (KLJ), Independent Research Fund Denmark 2025-00028B (GNH), Innovation Fund Denmark grant 9122-00012B (ATS), IMK Almene Fond (MDC, AMHE), Fulbright Denmark (EGP), and the European Union’s Horizon 2020 research and innovation programme under the Marie Sklodowska-Curie grant agreement No 814244 (ABIS, CH).

## Conflict of Interest

The lipidated dimeric peptides, their usage, and their extended utilization are disclosed in patent application WO2021/176094 currently being processed by patent authorities. KLM and ATS have ownership interests and are co-founders of Zyneyro, a company having exclusive license rights on the patent, which is owned by the University of Copenhagen, Denmark.

## Notes

### Summary of Updates

We have added data on the pain-relieving effect of mPD5 in the CFA and SNI model in female mice, which was previously only performed in male mice. We have considerably broadened the side-effect profiling and report no effect on acute nociception, learning and memory, or male fertility as well as no abuse liability. We have assessed the effect of mPD5 in PICK KO confirming PICK1 as the target for efficacy of mPD5. Extensive reassessment of the distribution of mPD5 following s.c. administration has been added to the manuscript, which strongly suggest that mPD5 does not distribute to the CNS but exerts its effect peripherally in the first order sensory neurons.

