## Supplemental information for "mPD5, a peripherally restricted PICK1 inhibitor for treating chronic pain"

### SUPPORTING INFORMATION

### Supplemental Table S1. SAXS data collection and fitting table.

| | | | | | |
|---|---|---|---|---|---|
| **Sample details** | mPD5 | | | | |
| Peptide | $C_{14}$-NPEG$_4$-(HWLKV)$_2$ | | | | |
| Buffer | 50mM Tris, 125mM NaCl, pH 7.4 | | | | |
| Extinction coefficient | 11 000 M$^{-1}$ cm$^{-1}$ | | | | |
| Molecular weight | 1874 g/mol | | | | |
| Peptide concentration | 0.19 - 9.37 mg/ml | | | | |
| **SAXS data collection details** | | | | | |
| Instrument | P12, Petra III, DESY [1] | | | | |
| Date for data collection | 09-05-2019 | | | | |
| Wavelength | 1.24 | | | | |
| Measured $q$-range (Å$^{-1}$) | 0.0025-0.73 | | | | |
| Absolute calibration | Water | | | | |
| Exposure time (ms) | 10 ms | | | | |
| Temperature (K) | 293.35 | | | | |
| **Software** | | | | | |
| Indirect Fourier transformations to obtain p($r$) | Bayersapp.org [2,3] | | | | |
| Data rebin | WillItRebin [4] | | | | |
| **Core Shell model fit** | | | | | |
| $N_{agg}$ | 18.7 | 21.5 | 22.2 | 21.0 | 19.0 |
| $v_{molec\text{-}hydrophilic}$ [$Å^3$] | 1985 | 1980 | 2012 | 2011 | 2012 |
| $v_{molec\text{-}hydrophobic}$ [$Å^3$] (fixed) | 377 | 377 | 377 | 377.1 | 377.1 |
| $\sigma_{relative}$ | 0.57 | 0.41 | 0.32 | 0.30 | 0.27 |
| $\chi^2$ | 1.1 | 1.5 | 2.5 | 14.5 | 15.1 |
| *Average model parameters* | | | | | |
| $N_{agg}$ | 20.5 | | | | |
| $v_{molec\text{-}hydrophilic}$ [$Å^3$] | 2012 | | | | |
| $v_{molec\text{-}hydrophobic}$ [$Å^3$] | 377 | | | | |
| $R_{total}$ [Å] | *22.6 (corresponding to a hydrophilic shell thickness of $D_{shell}$=10.4 Å)* | | | | |
| $R_{core}$ [Å] | *12.2* | | | | |

**Footnotes and references**

[1] C. E. Blanchet, A. Spilotros, F. Schwemmer, M. A. Graewert, A. Kikhney, C. M. Jeffries, D. Franke, D. Mark, R. Zengerle, F. Cipriani, S. Fiedler, M. Roessle, D. I. Svergun, *J Appl Crystallogr* **2015**, *48*, 431.
[2] S. Hansen, *Journal of Applied Crystallography* **2012**, *45*, 566.
[3] S. Hansen, *Journal of Applied Crystallography* **2014**, *47*, 1469.
[4] WillItFit MC Pedersen **2013.**

**Supplemental Table S2. Table of statistics**

| Figure | Test | Sample size | P-values |
|---|---|---|---|
| **1A-J** | NA | NA | NA |
| **2A-G** | NA | NA | NA |
| **3A** i.t. mPD5 administration in CFA-mice followed by von Frey | Two-way ANOVA followed by Dunnetts posthoc test of time 0 hrs versus 1, 5, 24 hrs | $n_{vehicle} = 5$<br>$n_{mPD5} = 6$ | mPD5:<br>$p_{1hr} = 0.0019$<br>$p_{5hrs} = 0.0296$<br>$p_{24hrs} > 0.9999$<br>Vehicle:<br>$p_{1hr} = 0.9969$<br>$p_{5hrs} > 0.9999$<br>$p_{24hrs} > 0.9999$ |
| **3B** s.c. mPD5 administration in CFA-mice followed by von Frey | Two-way ANOVA followed by Dunnetts posthoc test of time 0 hrs versus 1, 5, 24 hrs | $n_{saline\_saline} = 4$<br>$n_{CFA\_saline} = 5$<br>$n_{CFA\_mPD5} = 6$ | saline_saline:<br>$p_{1hr} = 0.4778$<br>$p_{5hrs} > 0.9999$<br>$p_{24hrs} = 0.9273$<br>CFA_saline:<br>$p_{1hr} = 0.9988$<br>$p_{5hrs} > 0.9999$<br>$p_{24hrs} = 0.9997$<br>CFA_mPD5:<br>$p_{1hr} < 0.0001$<br>$p_{5hrs} = 0.5171$<br>$p_{24hrs} = 0.9985$ |
| **3C** s.c. mPD5 (2, 10 or 50 μmol/kg) administration in CFA-mice followed by von Frey | Two-way ANOVA followed by Dunnetts posthoc test of time 0 hrs versus 1, 5, 24 hrs | $n_{50\ \mu mol/kg} = 5$<br>$n_{10\ \mu mol/kg} = 6$<br>$n_{2\ \mu mol/kg} = 5$ | 2 μmol/kg:<br>$p_{1hr} = 0.0476$<br>$p_{5hrs} = 0.9773$<br>$p_{24hrs} = 0.9773$<br>10 μmol/kg:<br>$p_{1hr} = 0.0426$<br>$p_{5hrs} = 0.1585$<br>$p_{24hrs} = 0.9998$<br>50 μmol/kg:<br>$p_{1hr} = 0.0001$<br>$p_{5hrs} = 0.0345$<br>$p_{24hrs} = 0.9690$ |
| **3D** | NA | NA | NA |
| **3E** s.c. mPD5 administration in CFA-mice followed by Hargreaves test | Two-way ANOVA followed by Dunnetts posthoc test of time 0 hrs versus 1, 5, 21 hrs | n = 6 in both groups | mPD5:<br>$p_{1hr} = 0.0001$<br>$p_{5hrs} = 0.3282$<br>$p_{21hrs} = 0.9620$<br>PBS:<br>$p_{1hr} = 0.8295$<br>$p_{5hrs} = 0.8130$<br>$p_{21hrs} > 0.9999$ |

| | | | |
|---|---|---|---|
| **3F** s.c. mPD5 administration in CFA-mice followed by marble burying test | one-way ANOVA with Tukey multiple comparison | n = 12 in each group | CFA:<br>p = 0.0024<br>mPD5:<br>p = 0.6802<br>PBS:<br>p = 0.0209 |
| **3G** s.c. mPD5 administration in CFA-mice followed by elevated plus maze | one-way ANOVA with Tukey multiple comparison | n = 12 in each group | CFA:<br>p = 0.0299<br>mPD5:<br>p = 0.6903 |
| **3H** NA | NA | NA | NA |
| **3I** s.c. mPD5 administration in CFA-mice followed by sePP | Unpaired t test | $n_{PBS}$= 15<br>$n_{mPD5}$ = 12 | mPD5vsPBS:<br>p = 0.0403 |
| **3J** s.c. mPD5 administration in control mice followed by sePP | Unpaired t test | n = 7 in both groups | mPD5vsPBS:<br>p = 0.9884 |
| **4A** s.c. mPD5 administration in SNI-mice followed by von Frey | Two-way ANOVA followed by Dunnetts posthoc analysis | n = 6 in each group | PBS:<br>$p_{1hr}$ = 0.8530<br>$p_{2hrs}$ = 0.9108<br>$p_{3hrs}$ = 0.9054<br>$p_{4hrs}$ = 0.7305<br>$p_{6hrs}$ = 0.6840<br>2 µmol/kg:<br>$p_{1hr}$ = 0.0633<br>$p_{2hrs}$ = 0.4194<br>$p_{3hrs}$ = 0.9423<br>$p_{4hrs}$ = 0.9998<br>$p_{6hrs}$ = 0.9997<br>10 µmol/kg:<br>$p_{1hr}$ < 0.0001<br>$p_{2hrs}$ < 0.0001<br>$p_{3hrs}$ < 0.0001<br>$p_{4hrs}$ = 0.0668<br>$p_{6hrs}$ = 0.9565 |
| **4B** i.p. mPD5 administration in STZ-mice followed by von Frey | Two-way ANOVA followed by Dunnetts posthoc test of time 0 hrs versus 1, 2, 4, 6 and 24 hrs | $n_{pregabalin}$ = 10<br>$n_{mPD5}$ = 10<br>$n_{PBS}$ = 9 | Pregabalin:<br>$p_{1hr}$ = 0.0046<br>$p_{2hrs}$ < 0.0001<br>$p_{4hrs}$ = 0.0046<br>$p_{6hrs}$ > 0.9999<br>Vehicle:<br>$p_{1hr}$ > 0.9999<br>$p_{2hrs}$ = 0.9961<br>$p_{4hrs}$ = 0.9961<br>$p_{6hrs}$ = 0.9998<br>$mPD5_{2µmol/kg}$:<br>$p_{1hr}$ = 0.0186<br>$p_{2hrs}$ = 0.2315<br>$p_{4hrs}$ = 0.2315 |

| | | | |
|---|---|---|---|
| | | | $p_{6hrs} = 0.9998$<br>$mPD5_{10\mu mol/kg}$:<br>$p_{1hr} < 0.0001$<br>$p_{2hrs} < 0.0001$<br>$p_{4hrs} = 0.0894$<br>$p_{6hrs} = 0.9998$ |
| **4C** s.c. mPD5 administration in CIBP mice followed by von Frey | Two-way ANOVA followed by Dunnetts posthoc test of pre versus post + baselines | $n_{morphine} = 10$<br>$n_{mPD5} = 11$<br>$n_{PBS} = 11$ | $p_{pre\text{-}postmPD5} = 0.6484$<br>$p_{pre\text{-}postmorphine} < 0.0001$<br>$p_{pre\text{-}postPBS} = 0.7914$ |
| **4D** Hot water tail immersion test | Two-way ANOVA followed by Sidaks multiple comparison | $n_{morphine} = 8$<br>$n_{mPD5} = 8$<br>$n_{PBS} = 8$ | Baseline vs 1 hour:<br>$P_{PBS} = 0.0907$<br>$P_{morphine} < 0.0001$<br>$P_{mPD5} = 0.9204$ |
| **4E** Capsaicin test | One-way ANOVA followed by Dunnets posthoc test | | $P_{Vehiclevsmorhpine} < 0.0001$<br>$P_{VehiclevsmPD5} = 0.6899$ |
| **5A NA** | NA | NA | NA |
| **5B** s.c. mPD5 administration in female SNI-mice followed by von Frey | Two-way ANOVA followed by Dunnetts posthoc analysis | n = 6 in each group | Vehicle:<br>$p_{1hr} = 0.8736$<br>$p_{5hrs} = 0.8993$<br>$p_{24hrs} > 0.9999$<br>$mPD5_{2\mu mol/kg}$:<br>$p_{1hr} = 0.0006$<br>$p_{5hrs} = 0.3634$<br>$p_{24hrs} = 0.9987$<br>$mPD5_{10\mu mol/kg}$:<br>$p_{1hr} = 0.0001$<br>$p_{5hrs} = 0.2270$<br>$p_{24hrs} = 0.5107$<br>$mPD5_{50\mu mol/kg}$:<br>$p_{1hr} = 0.0042$<br>$p_{5hrs} = 0.3685$<br>$p_{24hrs} = 0.9178$ |
| **5C** s.c. mPD5 administration in SNI-mice followed by marble burying test | one-way ANOVA with Dunnetts posthoc analysis | n = 12 in each group | $P_{SNIPBS\ vs\ control} = 0.0138$<br>$P_{SNImPD5\ vs\ mPD5} = 0.0035$<br>$P_{control\ vs\ SNImPD5} = 0.8142$ |
| **5D** s.c. mPD5 administration in SNI-mice followed by sePP | Unpaired t test | n = 11 in both groups | mPD5vsPBS:<br>p = 0.4190 |
| **5E** recordings of ultrasonic vocalizations in SNI mice | one-way ANOVA of baseline and 1 hour with Dunnetts posthoc analysis | n = 6 in both groups | Baseline:<br>$P_{control\ vs\ SNImorphine} = 0.0216$<br>$P_{control\ vs\ SNImPD5} = 0.0065$<br>Hour 1:<br>$P_{control\ vs\ SNImorphine} = 0.0282$ |

| | | | |
|---|---|---|---|
| | | | $P_{control\ vs\ SNImPD5}$ = 0.0393 |
| **5F** s.c. mPD5 administration in wildtype and PICK1 knockout SNI-mice followed by von Frey | Two-way ANOVA followed by Dunnetts posthoc analysis | n = 6 in both groups | Wildtype_PBS:<br>$p_{1hr}$ = 0.9043<br>$p_{5hrs}$ = 0.9043<br>Wildtype_mPD5:<br>$p_{1hr}$ = 0.0008<br>$p_{5hrs}$ = 0.0261<br>Knockout_PBS:<br>$p_{1hr}$ = 0.6818<br>$p_{5hrs}$ = 0.3568<br>Knockout_mPD5:<br>$p_{1hr}$ = 0.7598<br>$p_{5hrs}$ = 0.4684 |
| **6A** Open field locomotion of naïve mice in 5 min bins | | | |
| **6B** Total open field locomotion of naïve mice | | | |
| **6C** | NA | NA | NA |
| **6D** sperm count of males in the fertility study | | | |
| **6E** number of pubs per litter in the fertility study | | | |
| **6F** Drop in weight of females (per pub) in fertility study | | | |
| **6G** | NA | NA | NA |
| **6H** | NA | NA | NA |
| **6I** Latency to target on probe test on day 5 of Barnes' Maze test | | | |
| **6J** Total locomotion at probe test on day 5 of Barnes' Maze test | | | |
| **6K** Latency to target on probe test on day 9 of Barnes' Maze test | | | |
| **6L** Total locomotion at probe test on day 9 of Barnes' Maze test | | | |
| **6M** | NA | NA | NA |
| **6N** Locomotion during conditioning in the conditioned place preference test | | | |

| | | | |
|---|---|---|---|
| **6O** Posttest following 8 days of conditioning in the conditioned place preference test | | | |
| **7A** repeated s.c. mPD5 administration in SNI mice followed by von Frey | Two-way ANOVA followed by Dunnetts posthoc analysis compared to baseline post SNI | n = 8 in each group | <u>mPD5$_{2\mu mol/kg}$:</u><br>day 1<br>$p_{1hr} = 0.0105$<br>$p_{5hrs} = 0.1037$<br>$p_{23hrs} = 0.6065$<br>day 2<br>$p_{1hr} = 0.0681$<br>$p_{5hrs} = 0.2325$<br>$p_{24hrs} = 0.9996$<br><u>mPD5$_{10\mu mol/kg}$:</u><br>day 1<br>$p_{1hr} = 0.0077$<br>$p_{5hrs} = 0.2249$<br>$p_{23hrs} = 0.9794$<br>day 2<br>$p_{1hr} = 0.0082$<br>$p_{5hrs} = 0.0241$<br>$p_{24hrs} = 0.9997$<br><u>mPD5$_{50\mu mol/kg}$:</u><br>day 1<br>$p_{1hr} = 0.0026$<br>$p_{5hrs} = 0.0368$<br>$p_{23hrs} = 0.9806$<br>day 2<br>$p_{1hr} = 0.0008$<br>$p_{5hrs} =0.0112$<br>$p_{24hrs} = 0.7490$ |
| **7B** | NA | NA | <u>NA</u> |
| **7C** sustained s.c. mPD5 administration in SNI mice followed by von Frey | Two-way ANOVA with Dunnetts multiple comparison vs baseline before treatment | n = 8 in each group | <u>Group A:</u><br>$hour_{1-10}$ $p < 0.0001$<br>$hour_{11}$ $p = 0.0011$<br>$hour_{22}$ $p = 0.0121$<br>$hour_{23}$ $p = 0.1169$<br>$hour_{25}$ $p = 0.0023$<br>$hour_{27}$ $p = 0.1125$<br>$hour_{49}$ $p = 0.1331$<br>$hour_{121}$ $p = 0.9957$<br><u>Group B:</u><br>$hour_{1}$ $p < 0.0001$<br>$hour_{4.5}$ $p = 0.1927$<br>$hour_{6}$ $p = 0.0775$<br>$hour_{7}$ $p = 0.1063$<br>$hour_{8}$ $p = 0.7768$<br>$hour_{9}$ $p = 0.8516$<br>$hour_{10}$ $p = 0.9997$<br>$hour_{11}$ $p > 0.9999$ |

| | | | |
|---|---|---|---|
| | | | hour$_{22}$ p = 0.9994<br>hour$_{23}$ p > 0.9999<br>hour$_{25}$ p = 0.9997<br>hour$_{27}$ p > 0.9999<br>hour$_{49}$ p = 0.9996<br>hour$_{121}$ p = 0.9999 |
| **7D** | NA | NA | NA |
| **7E** | NA | NA | NA |
| **7F** s.c. saline/mPD5/aPD5/sPD5 administration in SNI mice followed by von Frey | Two-way ANOVA followed by Dunnetts posthoc analysis | n = 6 in each group | vehicle:<br>$p_{2hr} = 0.7257$<br>$p_{8hrs} = 0.6137$<br>$p_{13hrs} = 0.4137$<br>mPD5:<br>$p_{2hr} < 0.0001$<br>$p_{8hrs} = 0.8933$<br>$p_{13hrs} > 0.9999$<br>sPD5:<br>$p_{2hr} = 0.0365$<br>$p_{8hrs} = 0.9501$<br>$p_{13hrs} = 0.9472$<br>aPD5:<br>$p_{2hr} = 0.0058$<br>$p_{8hrs} = 0.4245$<br>$p_{13hrs} = 0.1971$ |

**Figure S1.**

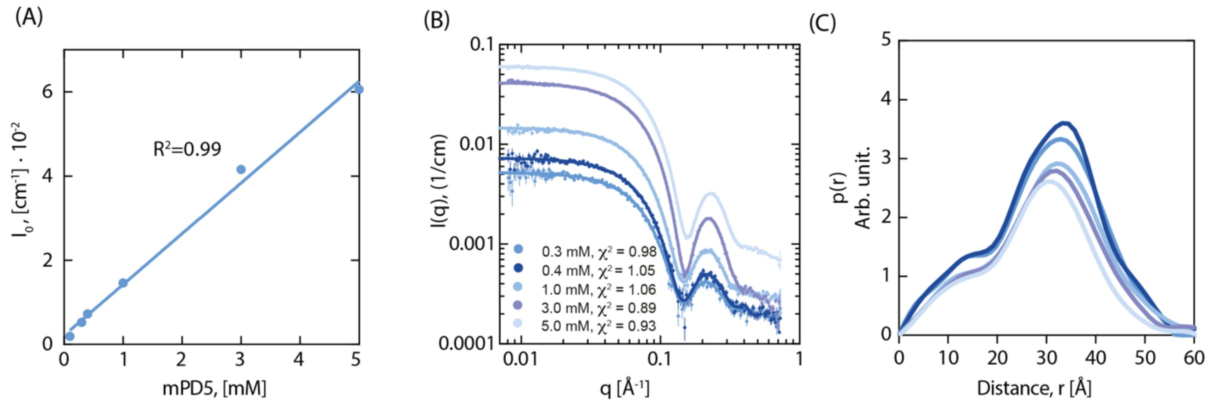

**Supplementary Figure S1. Biophysical characterization of mPD5. (A)** Plot of forward scattering ($I_0$) vs. concentration for increasing concentrations of mPD5. **(B)** The Indirect Fourier Transformation fits to the SAXS data (see Supplemental Table S1 for fitting parameters). **(C)** Pair distance distribution function (PDDF), of different concentrations (as indicated in (Figure S1B)) of mPD5.

**Figure S2.**

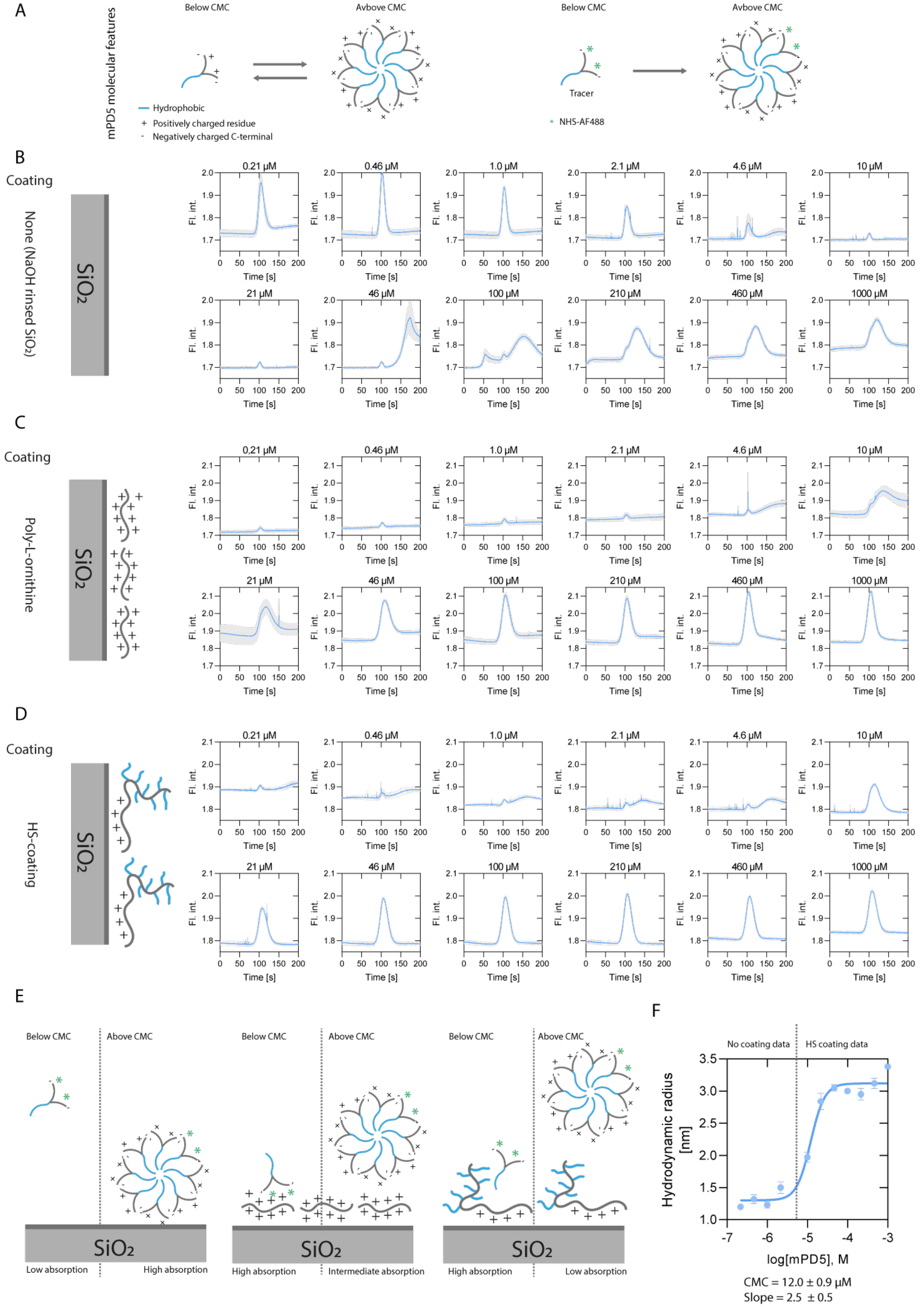
A
mPD5 molecular features
Below CMC
Avbove CMC
Below CMC
Avbove CMC
Tracer
Hydrophobic
Positively charged residue
Negatively charged C-terminal
NHS-AF488
B
Coating
None (NaOH rinsed SiO2)
SiO2
C
Coating
Poly-L-ornithine
D
Coating
HS-coating
Fl. int.
Time [s]
0.21 µM
0.46 µM
1.0 µM
2.1 µM
4.6 µM
10 µM
21 µM
46 µM
100 µM
210 µM
460 µM
1000 µM
E
Below CMC
Above CMC
Low absorption
High absorption
Intermediate absorption
F
No coating data
HS coating data
Hydrodynamic radius [nm]
log[mPD5], M
CMC = 12.0 ± 0.9 µM
Slope = 2.5 ± 0.5

**Supplementary Figure S2. FIDA analysis of mPD5. (A)** Molecular properties of mPD5. **(B)** Taylorgrams for different concentrations of mPD5 using an uncoated capillary. **(C)** Taylorgrams for different concentrations of mPD5 using a Poly-L-ornithine capillary. **(D)** Taylorgrams for different concentrations of mPD5 using an HS-coated capillary. **(E)** Graphical representation of mPD5 absorption using different coating strategies (B-D). **(F)** Binding isotherm of mPD5 combining optimal coatings for different concentrations results in a CMC of 12 μM. Data was plotted using GraphPad Prism 8.3, and raw data was fitted using FIDA Software (2.0), and resulting hydrodynamic radii were plotted and fitted using a four parameters saturation binding curve with variable slope.

**Figure S3**

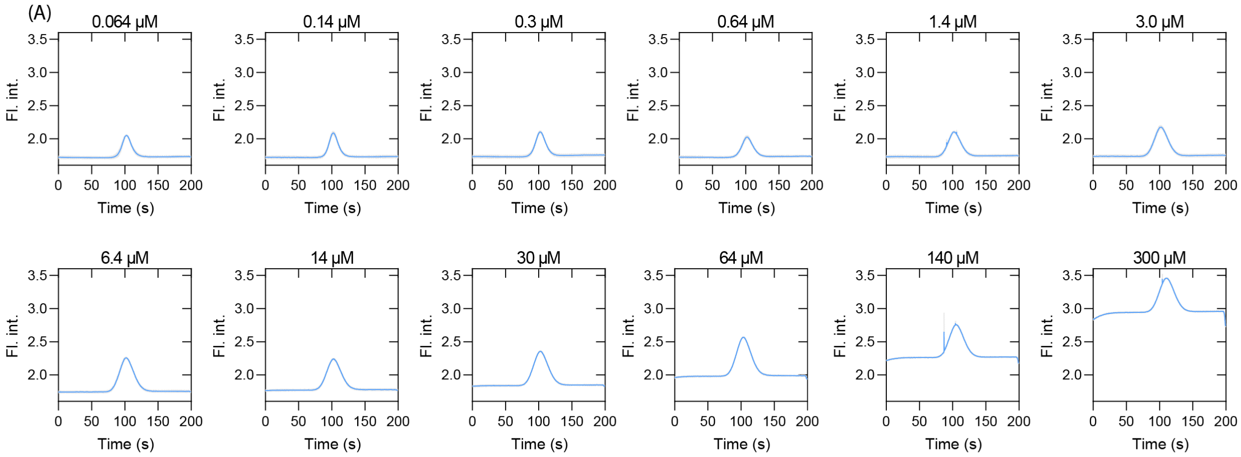

**Supplementary Figure S3 mPD5 binding to serum proteins. (A)** Taylorgrams from Flow-induced dispersion analysis (FIDA) for different concentrations of HSA and a fixed concentration of mPD5-AF488 (100 nM) using an HS-uncoated capillary.

**Figure S4**

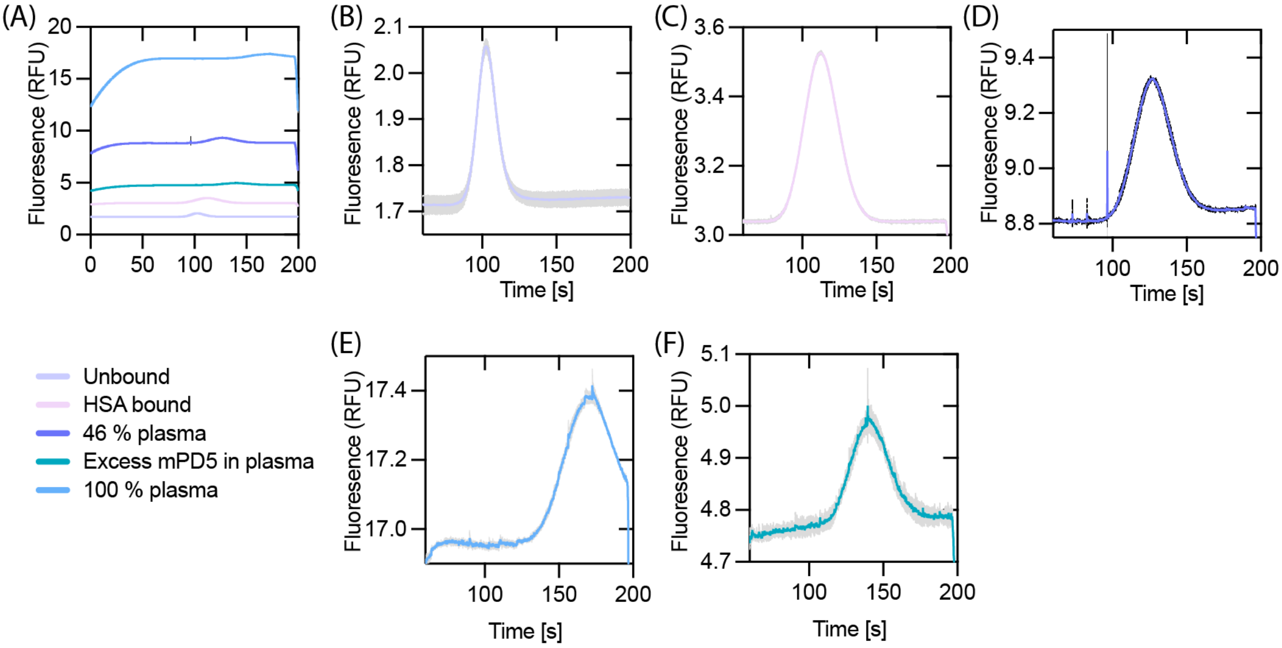

**Supplementary Figure S4 (A)** Taylorgrams for mPD5-AF488 in indicated conditions. **(B)** mPD5-AF488 (100 nM) in an unbound state. **(C)** mPD5-AF488 in the HSA bound state (see also Supplementary Figure S3, 300 μM HSA). **(D)** mPD5-AF488 in 46% human plasma. **(E)** mPD5-AF488 in 100% human plasma. **(F)** mPD5-AF488 in 25% supplemented with 21 mM mPD5 human plasma. n = 3 with errors as grey interval depicted as SEM.

**Figure S5.**

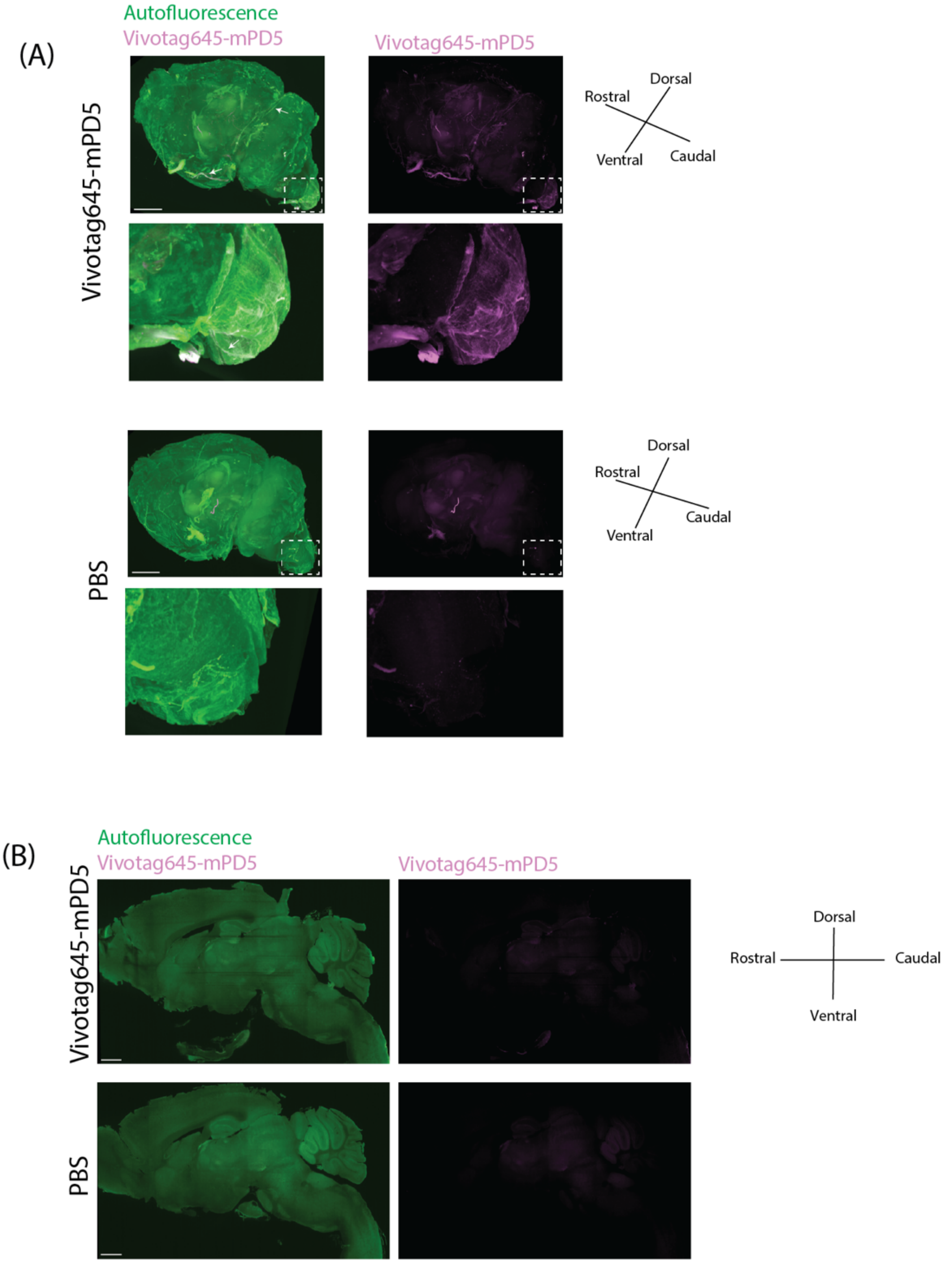

**Supplementary Figure S5. (A)** Maximum projection of 3D imaged cleared brain with Vivotag645-mPD5 (magenta) and autofluorescence (green). Scale bar = 2000 µm. Dashed boxes outline the magnified view of the most rostral part of the spinal cord. The arrows point to tissue structures with signal from Vivotag645-mPD5. **(B)** Optical section of 3D imaged brain in sagittal view with Vivotag645-mPD5 (magenta) and autofluorescence (green). Scale bar = 1000 μm.

**Figure S6.**

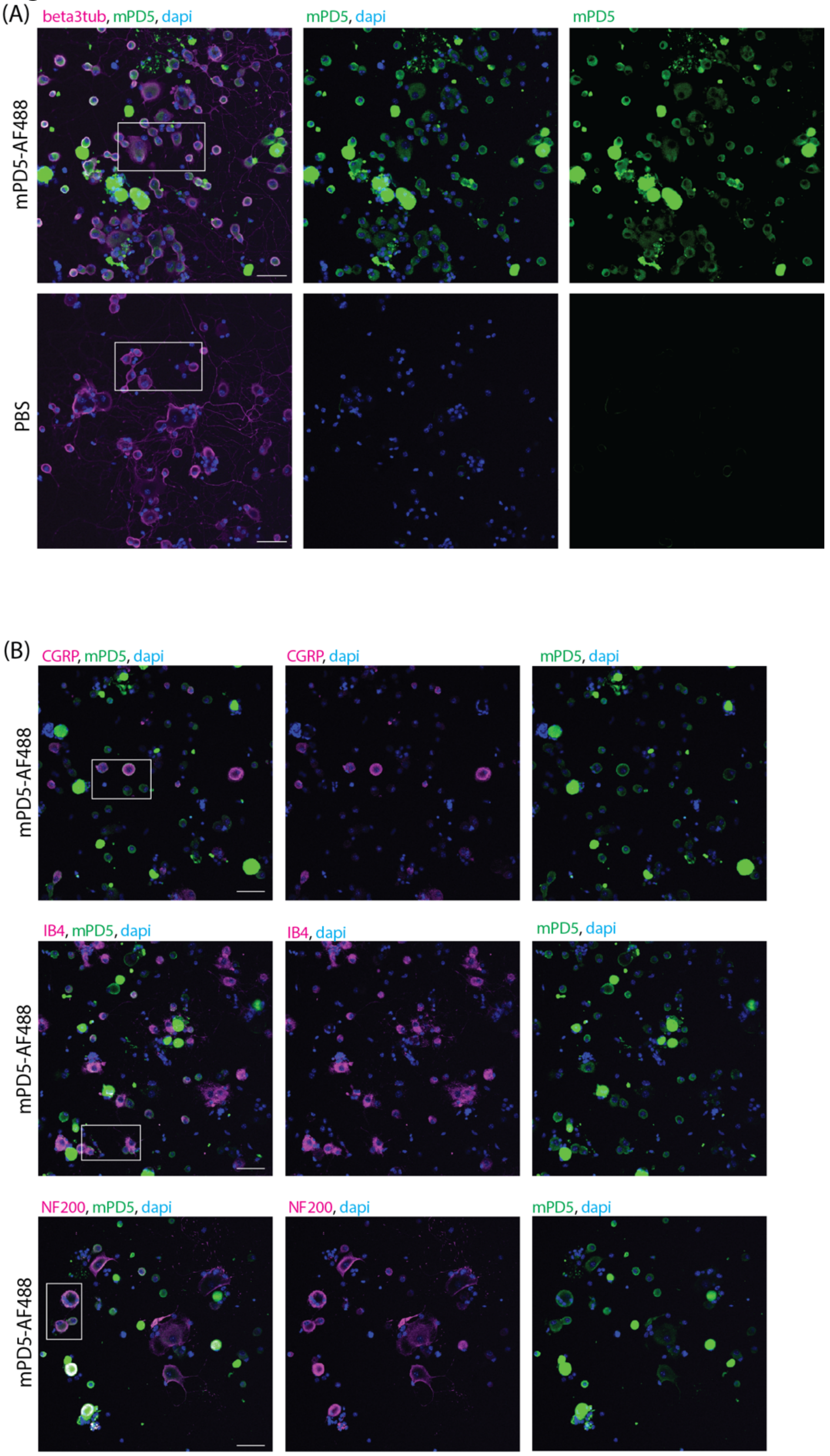

**Supplementary Figure S6. (A)** Primary dorsal root ganglia culture stained against neurons with beta 3 tubulin (magenta), mPD5-AF488 (green), and nuclei (blue). White boxes highlight the view shown in Figure 2A. Scale bar = 50 µm. **(B)** Primary dorsal root ganglia culture stained against neuronal subtype markers CGRP, IB4 or NF200 (magenta), mPD5-AF488 (green), and nuclei (blue). White boxes highlight the view shown in Figure 2B. Scale bar = 50 µm.

### Figure S7

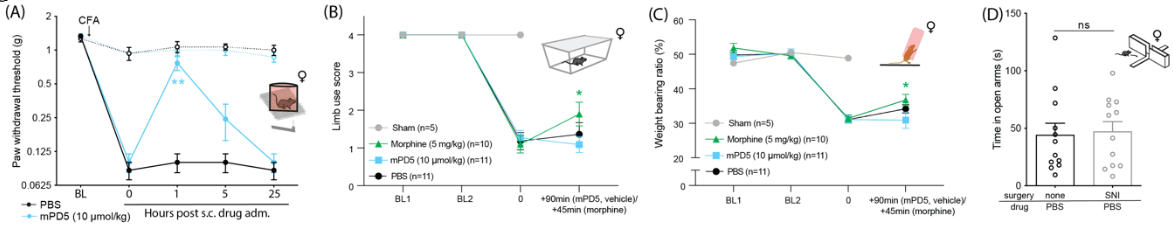

**Supplementary Figure S7. (A)** Paw withdrawal threshold (PWT) before and after induction of inflammatory pain (i.pl. CFA injection) and subcutaneous (s.c.) treatment with mPD5 (10 µmol/kg) or PBS. Two-way ANOVA with Dunnetts posthoc test. n = 6. Dashed line = contralateral paw. **(B+C)** Efficacy of mPD5 in mice with cancer induced bone pain or sham animals. $n_{morphine} = 10$, $n_{mPD5} = 11$ and $n_{PBS} = 11$. **(B)** Limb use score before and after induction of cancer induced bone pain (sham or NCTC 2472 cell inoculation) and subcutaneous (s.c.) treatment with mPD5, morphine or PBS. Wilcoxon matched-pairs signed rank test (non-parametric data). **(C)** As in panel B, but for weight bearing. Two-way ANOVA with Dunnetts posthoc test of pre versus post and baselines**.** All data is expressed as mean ±SEM. Abbreviations; adm. = administration, BL = baseline, Lu = limb use score, s.c. = subcutaneous, Wb =weight bearing ratio.

**Figure S8**

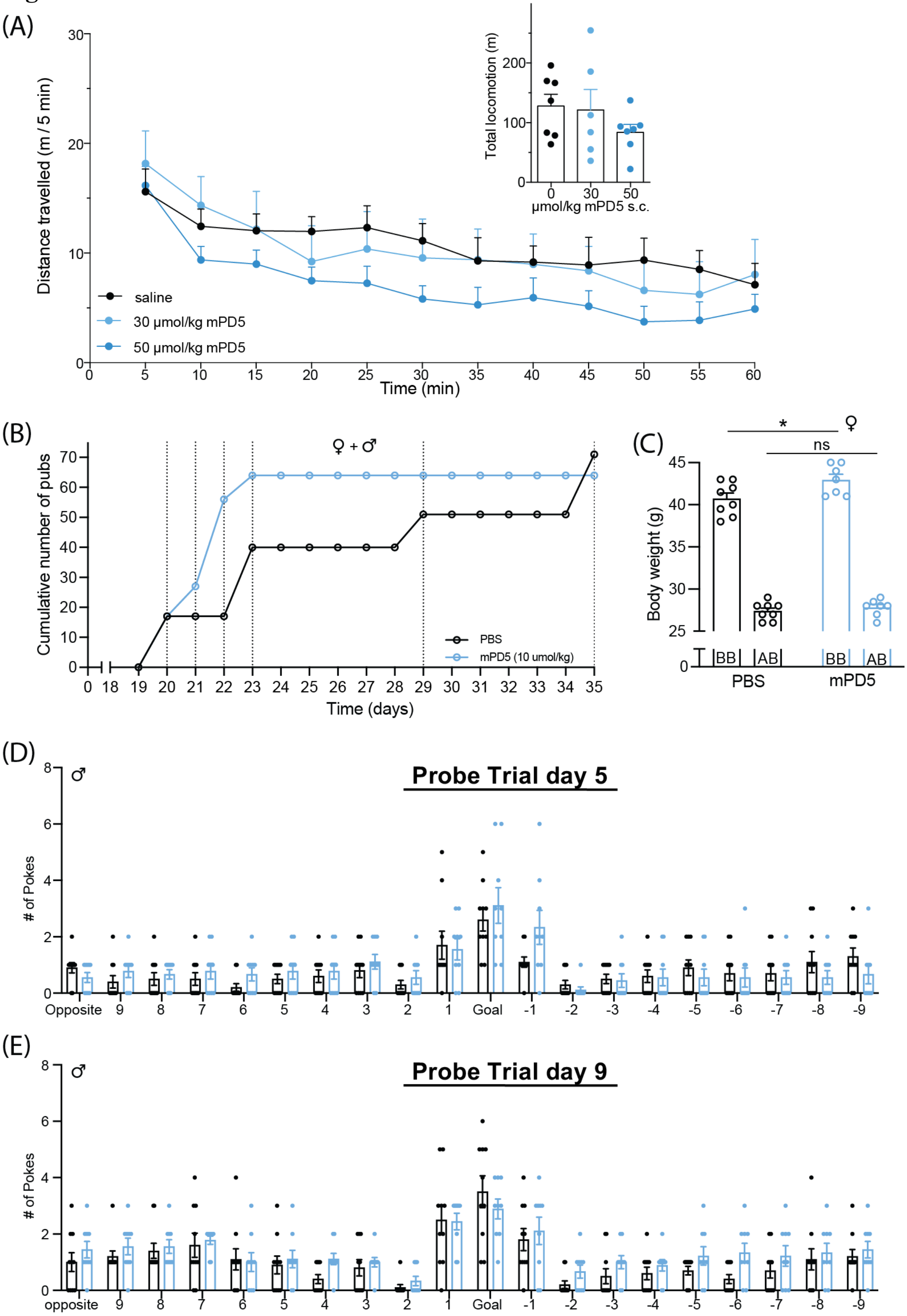

**Supplementary Figure S8. Effect of mPD5 on naïve animals.**

**(A)** One hour open field test of naïve mice 60 min after subcutaneous injection of saline or mPD5 with distance travelled (5 min bins) and total distance (insert). Two-way ANOVA with

Dunnetts posthoc analysis. $n_{saline} = 7$, $n_{30} = 6$, $n_{50} = 7$. **(B+C)** Supplementary data for figure 6C-F. **(B)** Cumulative number of pups of the two groups. **(C)** Body weight of the pregnant female mice before birth (BB) and after birth (AB) of the two groups. **(D+E)** Supplementary data for figure 6G-L. **(D)** Number of nose pokes into the 20 different holes of the Barnes Maze during the probe trial on day 5. **(E)** Number of nose pokes into the 20 different holes of the Barnes Maze during the probe trial on day 9. All data is expressed as mean ±SEM.

### METHODS

**Sex as a biological variable.**
Our study examined male and female mice, and similar findings are reported for both sexes.

**Peptides.**
All peptides were ordered from WuXI AppTec (*Shanghai, China*) or TAG-Copenhagen (*Copenhagen, Denmark*) with >95% purity, validated by MS and UPLC. D5 (DAT-C5, HWLKV), NPD5 ($NPEG_4$-$(HWLKV)_2$), TPD5 (YGRKKRRQRRR-$NPEG_4$-$(HWLKV)_2$), mPD5 (myristyol-$NPEG_4$-$(HWLKV)_2$). 5FAM-TD (5FAM-ahx-YGRKKRRQRRRHWLKV), 5FAM-D5 (5FAM-ahx-HWLKV).

**Assessment of solubility and stability of mPD5.**
Solubility was determined by visual inspection of samples dissolved in increasing concentration in 10 mM PBS as judged by full transparency and monophasic appearance. Stability was addressed by REDGLEAD for four concentrations (2, 20, 50, and 200 µM), by leaving mPD5 in PBS at 5 and 25 ºC for 30 days followed by HPLC-UV-MS.

**Protein expression and purification**.
Full length rat PICK1 (pET41) was, as described earlier (1), expressed in BL21-DE3-pLysS and grown at 37 ºC, induced at $OD_{600}$= 0.6 with 1 mM IPTG, and grown 16 hours at 20 ºC. Cultures were harvested and re-suspended in 50 mM Tris, 125 mM NaCl, 2 mM DTT (Sigma), 1% Triton X-100 (Sigma), 20 µg/mL DNAse 1 and half a tablet of cOmplete protease inhibitor cocktail (Roche) pr. 1 L culture. The resuspended pellets were frozen at -80 ºC for later purification. The lysate was cleared by centrifugation (36,000 x g for 30 min at 4 ºC), and the supernatant was incubated with Glutathione-Sepharose 4B beads (GE Healthcare) for two hours at 4 ºC under gentle rotation and then centrifuged at 4,000 x g for five min. The supernatant was removed, and the beads were washed twice in 35 mL 50 mM Tris, 125 mM NaCl, 2 mM DTT, and 0.01% Triton-X100. The beads were transferred to PD-10 Bio-Spin® Chromatography columns (Bio-Rad) and washed with an additional three column volumes. Each column was sealed and 0.075 U/µL, Novagen® was added for cleavage O/N at 4 ºC under gentle rotation. PICK1 was eluted on ice and concentration was measured on a NanoDrop 3000.

**Fluorescence polarization binding.**
The competition binding assay was carried out using a fixed concentration of PICK1 (0.5 µM) and fluorescent tracer (5 nM 5-FAM-TD5 or 20nM 5FAM-D5) incubated with increasing concentrations of unlabelled peptides (D5, TPD5, mPD5) using black half-area Corning non-binding surface 96 well plates (Sigma-Aldrich, Ref. no. 3686). The plates were incubated 20 min on ice and the fluorescence polarization was measured on an Omega POLARstar plate (BMG LABTECH) reader using excitation filter at 485 nm and long pass emission filter at 520 nm. The data was plotted and fitted to a competitive binding One Site fit using GraphPad Prism 8.3.

**Size Exclusion Chromatography (SEC).**
Recombinant PICK1 (40 µM) was mixed for at least 20 min with mPD5 (10 µM) and run on a Superdex200 Increase 10/300. Similarly, mPD5 and PD5 were dissolved in 50 mM Tris (pH 7.4), 125 mM NaCl, 0.01% Triton X-100, 2mM DTT and run on Superdex200 Increase 10/300 measuring the absorption at 254 nm and 280 nm, due to detector saturation at high mPD5 concentrations, data was plotted for 254 nm.

**Flow-induced dispersion analysis (FIDA).**

FIDA was carried out using a fluorescently labelled (AF488-NHS) mPD5 analogue, and fluorescence was recorded at 488 nm using mPD5-AF488 (25% labelling efficiency) as a tracer (100 nM), using the standard protocol recommended by the manufacturer, in short, mPD5 and mPD5-AF488 (25% labelling efficiency) was loaded to the FIDA1 instrument, and peptide sample was injected into the capillary followed by a buffer injection. The diffusion of the peptide could then be observed using 488 nm fluorescence, and the hydrodynamic radius was calculated using the FIDA software 2.0 using a single Gaussian distribution fit, at 75% and curve smoothing. Resulting hydrodynamic radius was plotted using GraphPad Prism 8.3.

**Small Angle X-ray Scattering (SAXS).**

Concentration series of mPD5 ranging from 0.18 mg/mL to 9.37 mg/mL was prepared in buffer containing 50 mM Tris-HCl (pH 7.4) and 125 mM NaCl. Samples were measured at the P12 SAXS Beamline, PetraIII, DESY, Hamburg Germany (2). Initial data processing including radial averaging and conversion of the data into absolute scaled scattering intensity, I(q), as a function of the scattering vector q, where q= 4π sin(θ)/λ (θ = half scattering angle, λ = x-ray wavelength) were done using the standard software, procedures and settings at the beamline (2). The processed scattering data was merged, buffer subtracted, and binned using WillItRebin with a binning factor of 1.02. The modelling of the SAXS data was performed using a molecular constrained core-shell model for polydisperse spherical micelles. In brief, it was assumed that the peptide aggregated into polydisperse spherical micelles. Here, the hydrophobic tails form the core, and these are surrounded by the hydrophilic part (the shell) in a spherical micelle. For the modelling, total scattering lengths of 2.26e-10 cm and 2.96e-11 cm were used for the headgroups and tail, respectively. These were calculated by multiplying the number of electrons in the PD5 headgroup (801 electrons) and the $C_{13}$ alkyl chain (105 electrons) by the scattering length of the electron which is just the classical electron radium of 2.82 fm. For the scattering length density calculations, it was assumed that the molecular volume of the $C_{13}$ alkyl chain, $v_{molec\text{-}hydrophobic}$, was 377 Å$^3$ as estimated by Tanfords empirical formula ($V = 27.4 + 26.9n_C$) where $n_C$ denotes the number of carbons. i.e. 13 in this case (3). The hydrophilic headgroup, $v_{molec\text{-}hydrophilic}$, was as a first estimate assumed to have a mass density corresponding to that of a soluble protein (i.e. 1.35 g/cm$^3$), this yielded a molecular volume of 2078 Å$^3$. This value was then taken as a free parameter in the fits and refined to a molecular volume of ~2010 Å$^3$ corresponding to a mass density of 1.39 g/cm$^3$ and with very little variation over the fits to the different sample concentrations. Excess scattering length densities of the core and shell of the micelle were then calculated in relation to the scattering length density of water (9.4e10 1/cm$^2$). Using this molecular constrained model, the volumes of the core and of the shell were coupled internally through the fitted aggregation number, $N_{agg}$ and the implicit assumption of one PD5 headgroup per one $C_{13}$ alkyl chain. The relative polydispersity, $\sigma_{relative}$ was described by a Gauss function of $N_{agg}$. The Gauss was truncated at ±3σ. Through the fits, this value is refined to $\sigma_{relative}$ values of the aggregation number at ~30% for the highest concentrations to up to 50% for the lowest concentrations. Note, however, that this variation in the particle volume translates to about 10%-15% variation of the particle radius. A full overview of the resulting fit-parameters is provided in Supplemental Table S1. As usual a small surface roughness (typical at the order of 2Å) and a small constant background were also necessary for the model to converge (4). A more thorough description of the principle of molecular constrained modelling of simple micelle systems can be found in (5). The model module "PolydisperseMicelles" from the WillItFit software was used for the fitting (4).

**Animals.**

Unless otherwise specified, wild type C57BL/6NRj mice (male and female, aged eight-weeks at the beginning of experiment) were used (Janvier, France). Animals were allowed at least seven days of habituation to the animal facility before initiation of experiment. Mice were group-housed in a temperature-controlled room, maintained on a 12/12hour light/dark cycle (lights on at 6 A.M) with free access to standard rodent chow (Altromin 1342 (Brogaarden, Denmark)) and water.

**Drug administration and dosing for animal experiments.**
All compounds were diluted in PBS or saline (depending on control) and administered through different routes i.t. (7 µL/mouse)), i.p. (10 µL/g) or s.c. (10 µL/g)) as indicated. Doses of mPD5 were based on previous studies with TPD5 (6) with a subcutaneous dose-range included in the CFA studies further guiding dose selection for SNI, STZ and CIBP studies. Pregabalin dose was chosen based on previous experience from Phenotype Expertise (Marseilles, France). In the CIBP model, a dose of 5 mg/kg morphine was chosen to avoid hyperactivity, though known to provide only partial pain relief in this model. In remaining experiments, 5 mg/kg morphine was chosen, known to provide full pain relief, except for the conditioned place preference experiment, where 10 mg/kg was used instead in accordance with literature.

**Tissue clearing:**
C57BL/6 mice were injected s.c. with 10 umol/kg of mPD5 conjugated to Vivotag645 (WuXi, China) or PBS one hour prior to transcardial perfusion with PBS followed by 4% PFA (n=3). The spinal columns and the brains were dissected and post fixated in 4% PFA overnight at 4 °C. Next, the samples were washed in PBS with 0.02% sodium azide and cleared with a protocol combining elements from PEGASOS (7) and iDISCO (8) which obtained sufficient transparency while still keeping the Vivotag645 fluorophore intact. For the spinal columns, bone was decalcified with 20% EDTA tetrasodium salt (Sigma-Aldrich) in demineralized water for 4 days at 37 °C. Both spinal columns and brains were gently bleached with 25% Quadrol (Sigma-Aldrich) in demineralized water at 37 °C for 3 days followed by 5% ammonium (VWR, Denmark) in demineralized water for one day at 37 °C. Next, the samples were dehydrated in a dilution series of methanol (VWR, Denmark) in demineralized water (20%, 40%, 60%, 80%, 100%, and 100%) with 1.5 hours in each methanol concentration. The samples were delipidated overnight in a solution containing 2/3 dichloromethane (Sigma-Aldrich) and 1/3 methanol. The samples were washed two times in dichloromethane before index matching with ethyl cinnamate (Sigma-Aldrich). The spinal columns were imaged on a 3i Cleared Tissue LightSheet microscope with Slidebook software (3i). The Slidebook software was furthermore used to stitch the images. High resolution light sheet of the dorsal root ganglions (DRGs) and the imaging of the brains were performed with the Zeiss Lightsheet 7 with Zeiss Zen Black software. A custom Python script was used to stitch the imaged brains. After stitching visualization was optimized in Imaris software (Imaris Oxford instruments).

**Immunocytochemistry on primary dorsal root ganglion (DRG) neurons**
DRG cell cultures were prepared from adult C57BL/6 mice from Janvier. After transcardial perfusion with PBS, approximately 35 DRGs from one mouse were dissected and placed in ice cold Ham's F12 medium (Gibco). The tissue was incubated in enzyme mix (3 mL of 3 mg/mL dispase II from Sigma Aldrich and 200 µl of 12.5 mg/mL *clostridium histolyticum* collagenase from Sigma Aldrich) for one hour at 37 °C. The enzyme mix was removed and warm Ham's F12 medium was added. The cells were mechanically separated with serial trituration using fire-polished glass pipettes until the tissue was dissolved. The cells were passed through a 70 µm cell strainer (Sigma-Aldrich) and resuspended in 2 mL fresh F12 medium. Next, a 15% BSA (Sigma-Aldrich) solution was prepared, and the cells were gently placed on top without

mixing the solutions. After centrifugation at 1000 rpm for 10 min with acceleration = 1 and deceleration = 1 (Thermo Scientific, Heraeus Multifuge x3R with 75003180 rotor) the neurons were enriched in the pellet while myelin and some non-neuronal cells were in the interface between the 15% BSA solution and the Ham's F12 medium. The pellet was resuspended in cell medium (Ham's F12 with 10% FCS and 1% penicillin/streptomycin from Sigma) and plated on 13 mm Ø glass coverslips in 4 well trays. Previously to plating of the cells, the coverslips were coated over night at 4 °C with Poly-D-lysine 100 µg/ml (Gibco) followed by incubation of a 40 µl droplet of laminin 10 µg/ml (Sigma) for at least one hour at 37 °C. The laminin was removed just before plating the cells ensuring that the cells were restricted to the center of the coverslip. After letting the cells attach for 30 min in the cell incubator (37 °C, 5% $CO_2$) 500 µl cell medium was added to each well. The following day the cultures were incubated with 10 uM mPD5 conjugated to Alexa Fluor 488 (mPD5-AF488) (25% labelling efficiency) in Ham's F12 medium for one hour at 37 °C. The cells were gently washed in PBS before fixation at room temperature for 20 min with 4% PFA. Cells were permeabilized and blocked in PBS with 0.15% triton X-100 (Sigma) and 10% Donkey serum (Sigma-Aldrich) for one hour at room temperature. Next, cells were incubated with primary antibodies (see Supplemental Table S3 diluted in PBS with 0.15% triton and 10% donkey serum overnight at 4 °C protected from light. After 3x 5 min washes in PBS, the cells were incubated with secondary antibodies (see Supplemental Table S3) and IB4-647 (1:100, I32450, Invitrogen) diluted in PBS with 0.15% triton and 10% donkey serum for one hour at room temperature. The cells were washed 3x 5 min in PBS before mounting on a glass slide with DAPI Fluoromount-G (SouthernBiotech). The cells were imaged on a Zeiss confocal LSM780 microscope with a 20x objective and Zeiss Zen Black software.

**Supplemental Table S3: Primary and secondary antibodies used for DRG cultures.**

| Antibody | Dilution | Cat number | Supplier |
|---|---|---|---|
| Goat α-CGRP | 1:400 | Ab36001 | Abcam |
| Chicken α-NF200 | 1:500 | AB5539 | Millipore |
| Rabbit α-βIII tubulin | 1:1000 | Ab18207 | Abcam |
| Donkey α-goat 647 | 1:500 | A21447 | Invitrogen |
| Goat α-chicken 647 | 1:500 | A21449 | Life Technologies |
| Donkey α-rabbit 568 | 1:500 | A10042 | Invitrogen |

**Assessment of plasma levels and biodistribution of mPD5**.

The mPD5 plasma exposure was determined by (WuXI AppTec (*Shanghai, China*)) and done following s.c. injection of three male C57BL/6N (vendor: VTLH) mice (7-9 weeks old, fasted) with each concentration of mPD5 (2, 10 and 50 µmol/kg) in sterile PBS. Blood samples were taken 30 min, 1 hour, 2 hours, 5 hours, and 12 hours following injection. For the repeated administration experiment, the initial s.c. injection of 10 µmol/kg was followed by 4 s.c. administrations of 2 µmol/kg with one hour between injections. Following the last injection, blood samples (0.03 mL from saphenous vein or other suitable site) were collected in EDTA-K2 tubes at 5 min, 30 min, 1 hour, 5 hours, 12 hours, 24 hours, and 48 hours and centrifuged (app 4 °C, 3200 g, 10 min) for collection of plasma. Plasma was transferred to 96 well plates and quick-frozen over dry ice and kept at -60 °C until LC-MS/MS analysis. For LC-MS/MS analysis plasma with 0.25% Triton X100 in 50% MeOH/water (6 µL) was quenched with 160 µL of IS1 (internal standard in MeOH (Labetalol & tolbutamide & Verapamil & dexamethasone & glyburide & Celecoxib 100 ng/mL for each)) and the mixture was vortexed for 10 min at 800 rpm and centrifuged for 15 min at 3220 × g at 4 °C. An aliquot of 65 µL supernatant was transferred to a clean 96-well plate and centrifuged for 5 min at 3220 × g at 4 °C, then the supernatant was directly injected for LC-MS/MS analysis. Plasma half-lives were determined from three points in the elimination phase. In a separate experiment, biodistribution was assessed (plasma, CSF, spinal cord, whole brain) at two hours after single dose of 10 µmol/kg or one hour after last injection in the repeated administration paradigm (10 µmol/kg followed by four s.c. administrations of 2 µmol/kg with one hour between). At these times, plasma level were similar between the two paradigms. Plasma, CSF, and tissue samples (kept at 4 °C) were processed and analysed as described above.

**Locomotion.**
Concentration 10 µmol/kg: Wild type mice were habituated to the experimental room for a minimum of 60 min before initiation of the experiment. Female and male mice were run on separate days. Mice were injected s.c. with 10 µmol/kg mPD5 or sterile PBS directly before being placed in open field boxes (40 cm x 40 cm x 80 cm) for 2.5 hours with up to four mice in boxes at any one time. A video camera placed above the open field recorded the animals' behaviour and Ethovision XT 13 (Noldus) was used to track the distance travelled by the animal.
Concentration 30+50 µmol/kg: Animals were habituated to the experimental room for a minimum of 60 min before initiation of the experiment. At initiation of experiment, the mice were injected subcutaneously (s.c.) with mPD5 (30 or 50 µmol/kg, pH adjusted to approximately 7 with 5 M NaOH) or 0.9 % saline (B.Braun, Germany) and left in their home cage for 60 min. After exactly one hour, the mice were placed in an open field (40 cm x 40 cm x 80 cm) for 60 min. A video camera placed above the open field recorded the animals' behaviour and Ethovision XT 13 (Noldus) was used to track the distance travelled by the animal. The experiment was run in cohort with Figure 2B+C of (9)).

**Induction of inflammatory pain (CFA model).**
Injury was induced on the right hind paw, whereas the contralateral left hind paw was used as internal control of the C57BL/6J male and female mice (Janvier, France). The inflammatory pain was induced by injection of 50 µL undiluted Complete Freund's Adjuvant (CFA) (F5881, Sigma) unilaterally into the intraplantar surface of the right hind paw, whereas control mice were injected with the same amount of 0.9 % saline (B. Braun, Germany). All intraplantar injections were performed with an insulin needle (0.3 mL BD Micro-Fine) while the animal was under isoflurane anaesthesia (2%) for maximum 60 seconds. Von Frey was applied up to 11 days after unilateral CFA injection depending on the experiment. (For Figure 3A, the experiment was run in cohort with Figure 1A of (9)).

**Induction of neuropathic pain (SNI model).**
The initial SNI pain experiment (Figure 4A) was performed by Phenotype Expertise, Marseilles, France – Pain and CNS behaviour CRO under supervision of Stéphane Gaillard, PhD (CEO). Surgery was performed on 8-week-old C57BL/6J male mice (Charles River). The spared nerve injury surgery was performed on anaesthetized mice with a buprenorphine solution (0.05 mg/kg) injected s.c. 30 min before surgery (10 µL/g). For surgery, mice were anaesthetised with ketamine (100 mg/kg, i.p.) and xylazine (10 mg/kg, i.p.) (volume 10 µL/g)

with 100 μL of 1% xylocaine solution infiltrated at the site of the incision. The distal trifurcation of the sciatic nerve was identified, and the tibial and common peroneal branches were ligated with polypropylene nonabsorbable 6-0 sutures (Ethicon) and 1 mm was cut out, leaving the sural branch intact. The wound was closed with sutures, and the animals were allowed to recover and returned to their cages. Two subcutaneous injections of buprenorphine (0.05 mg/kg) were performed with an 8 to 12 hours delay decreasing pain due to the surgery act. seven days post-surgery, a decrease of threshold response to von Frey filaments of ipsilateral hind-paw was confirmed by von Frey filaments, corresponding to neuropathic pain condition. The mechanical threshold response of the operated mice was measured with calibrated von Frey filaments ("up/down" method) and the 50% threshold (g) was calculated.
The remaining SNI pain experiments were performed in-house using the following procedure. SNI surgery was performed on the left hindleg of C57BL/6J male and female mice (Janvier, France) under 2% isoflurane anaesthesia. Skin on the lateral surface was incised between hip and knee followed by lengthwise division of the biceps femoris muscle leading to exposure of the three branches of the sciatic nerve. Sural branch was left intact, while the peroneal and tibial branches were ligated with a single surgical knot and axotomized distally of the ligation. Wounds were closed with surgical glue or medical clips and animals were monitored daily for signs of stress or discomfort, but in all cases recovered uneventfully.

**Induction of Diabetic neuropathy (STZ model).**
Diabetes was induced by a single i.p. injection of 200 μg/mL streptozocin (STZ) solution (100 μL/10 g, Sigma-Aldrich S0130, batch #WXBB7152V) in male seven-week-old C57BL/6J (Charles River) mice. Glycemia was tested before, and seven days after injection. All injected mice had blood glucose concentration >350 mg/dL from day seven and were used for analgesic testing of the compounds from day 14. One mouse had to be euthanized at seven days post injection. This experiment was performed by Phenotype Expertise, Marseilles, France - Pain and CNS behaviour CRO under supervision of Stéphane Gaillard, PhD (CEO).

**Induction of cancer induced bone pain (CIBP model).**
Osteosarcoma cell line NCTC 2472 (American Type Culture Collection, CCL-11TM) was cultured for two weeks prior to surgery in NCTC-135 medium (Sigma-Aldrich, Denmark) supplemented with sodium hydrogen carbonate and 10% equine serum (Sigma-Aldrich, Denmark). On the day of surgery, cells were washed with PBS and incubated with 3 mL trypsin-EDTA for five min followed by centrifugation (2800 rpm, 3 min). Supernatant was removed, and cell pellet resuspended in PBS to obtain a cell density of $10^7$ cells/mL.
Two separate cohorts of female C3H/HeNHsd mice (Envigo, Netherlands) of 7-8 weeks, were anaesthetized with 6.25 mg/kg xylazine (Rompun vet, 20 mg/mL, Bayer, KVP Pharma + Veterinär Produkte GmbH, Germany, Batch: KV02BHF) and 43.75 mg/kg ketamine (Ketaminol vet, 100 mg/mL, MSD Animal Health, AN Boxmeer, The Netherlands, Batch: A140A01) and received 1.0-1.2% isoflurane (1000 mg/g isoflurane, Attane vet, ScanVet, Piramal Critical Care Ltd., UK, Batch: G123J18B) during surgery. The mouse was placed on a 37˚C heating pad. Both hind legs were shaved, and a small horizontal incision was made under the knee followed by a vertical incision of approximately 0.5 cm until the patellar tendon was exposed. The patellar tendon was moved to the lateral side exposing the distal epiphysis of the femur. A hole was made in the middle of the distal epiphysis by gently pushing a 30 G needle (0.3x12 mm, 30 Gx1/2", Luer-Lock, CHIRANA T. Injecta, CH30012) through the bone while rotating. The position of the needle was confirmed by X-ray imaging (IVIS Lumina XR Apparatus, Caliper Life Sciences, Belgium). Following resuspension, 10 μL, corresponding to $10^5$ NCTC 2472 cells or PBS (sham mice), was injected in the bone marrow cavity with an insulin syringe (1 mL Luer, BD PlastipakTM, 303172) attached to a 30 G needle and left in the

femur for one min. Bone wax (Harvard Apparatus, BS4 59-9864) was immediately pushed into the hole with a curette until bleeding stopped. The skin wound was closed with two medical clips (Michel Suture Clips, 7.5x1.75 mm, Angthos, 12040-01). Post-surgical analgesia included s.c. injection of 5 mg/kg carprofen (Carprosan vet, 50 mg/mL mixed 1:25 in sterile ddH2O, Dechra, Le Vet Beheer B.V., Netherlands, Batch: 050520) right after surgery and the following day. Post-surgical care included daily severity scoring for four days, and medical clips removed. The test battery of the CIBP model was carried out 45 or 90 min after morphine and mPD5 administration respectively to accommodate the peak efficacy of the drugs.

**Mechanically evoked pain (von Frey test).**
Unless otherwise specified, von Frey occurred as described in the following. Animals were habituated to the experimental room for a minimum of 60 min before initiation of the experiment. Mechanical paw withdrawal threshold (PWT) was determined by von Frey measurements of both hind paws. Von Frey filaments ranging from 0.04 to 2 g (g = gram-forces) (0.04, 0.07, 0.16, 0.4, 0.6, 1.0, 1.4, 2.0) were used for determination of the PWT. Filaments in ascending order were applied to the frontocentral plantar surface of the hind paws. Mice were placed in PVC plastic boxes (11.5 cm x 14 cm) on a wire mesh and allowed minimum 20 min habituation prior of experiment initiation. Each von Frey hair was applied five times with adequate resting periods between each application and number of withdrawals recorded. The withdrawal threshold was determined as the von Frey filament eliciting at least three positive trials out of the five applications in two consecutive filaments. A positive trial was defined as sudden paw withdrawal, flinching and/or paw licking induced by the filament.
For the CIBP model, two habituation sessions were performed on separated days with mice placed in the von Frey equipment for 1-1.5 hours in the presence of the experimenter on day one and random von Frey pocking with the 0.6 g filament on day two. Von Frey monofilaments (North Coast Touch Test, North Coast Medical, Inc.) with target forces of 0.04g, 0.07g, 0.16g, 0.40g, 0.60g, 1.00g, 1.40g, 2.00g, and 4.00g were used. 50 % PWT was found using the "up-down method" ($PWT{=}10^{Xf+k\delta}$) starting with the 0.6 g filament. Positive response was defined as; withdrawal, shaking or grooming of paw following a 2-3 second stimulation. In case, of unclear response, stimulus was repeated after 10-15 seconds.
For the STZ model, the mechanical threshold response of the injected mice was measured with calibrated von Frey filaments ("up/down" method) and the 50% threshold (g) was calculated. 13 days post injection, a decrease of threshold response to von Frey filaments of the ipsilateral hind paw was confirmed by von Frey filaments, corresponding to diabetic neuropathy. After injection of test compounds on day 14, mechanical threshold was measured at 1 hour, 2 hours, 4 hours and 6 hours and again on day 15. Compounds were diluted in PBS (vehicle) and administered s.c. at 10 μL/g, in doses as indicated (gabapentin, mPD5). This experiment was performed by Phenotype Expertise, Marseilles, France - Pain and CNS behaviour CRO under supervision of Stéphane Gaillard, PhD (CEO).

**Thermally evoked pain (Hargreaves test).**
A baseline Hargreaves test was performed on day 0 before inflammatory pain was induced as described in section 2.13. In the Hargreaves test, the mouse was placed in a container on a glass table and left to habituate for at least 15 min. Using the Hargreaves apparatus, a heat beam from beneath was directed towards the hind paws of the mouse, and the paw withdrawal time was registered automatically. The cut off was set to maximum 30 seconds to avoid skin injury and the experiment was repeated 5 times for each paw with at least five min in between measurements. On days -1, 2, and 3, mice were habituated to the Hargreaves equipment by placing them in the boxes for about 30 min without directing the beam at them.

**Repeated and sustained administration of mPD5 in the SNI model.**
For the experiment with repeated administration of mPD5 (Figure 7A), male C57BL/6J (Janvier, France) mice underwent surgery leading to partial nerve injury, by cutting the peroneal and tibial nerves, producing hypersensitivity of the remaining sural nerve (SNI) as described in the "**Induction of neuropathic pain (SNI model).**"-section. On day 21 after surgery, hyperalgesia was confirmed by using von Frey filaments to induce mechanically evoked hypersensitivity and measure paw withdrawal threshold of the animals as described in the "**Mechanically-evoked pain (von Frey).**"-section. On the same day, mice were injected s.c. with 2, 10, or 50 µmol/kg mPD5 (10 µL/g) and evoked pain was tested again with the use of von Frey filaments at 1 and 5 hours after injection. 24 hours after the first injection, mice were injected s.c. with 2, 10, or 50 µmol/kg mPD5 (10 µL/g) again and evoked pain was tested with the use of von Frey filaments at 1, 5, and 24 hours after the second injection.
For the first experiment with sustained administration of mPD5 (Figure 7C), the mice underwent a baseline von Frey reading as described in the "**Mechanically-evoked pain (von Frey).**"-section and SNI surgery as described in "**Induction of neuropathic pain (SNI model).**"-section. Approximately 18 weeks (125 days) after SNI surgery, hyperalgesia was confirmed with von Frey. On the same day, mice were injected s.c. with 10 µmol/kg mPD5 and evoked pain was tested again at 1 hour after injection. Then, mice were injected s.c. with either 2+2+2 µmol/kg mPD5 (group A) or PBS (group B) once an hour for 3 hours. At 4,5 hours after the 10 µmol/kg mPD5 injection paw withdrawal threshold was measured again followed by a s.c. injection of 2 µmol/kg mPD5 in all animals. The PWT was measured again at 6 hours +7 hours + 8 hours + 9 hours + 10 hours + 11 hours + 22 hours + 23 hours + 25 hours + 27 hours + 49 hours + 121 hours after the initial 10 µmol/kg mPD5 injection performed on day 125.

**Marble burying test.**
Animals were habituated to the experimental room for a minimum of 60 min before initiation of the experiment. Mice were injected with saline, PBS or 10 µmol/kg mPD5 (10 µL/g) and left in their home-cage. At 55 min post injection, the mice were placed in a cage with approximately five cm of bedding and left to habituate for five min. One hour after injection, the bedding was rearranged to be level again, and 20 marbles were evenly spaced on top of the bedding. Mice were given 20 min to bury marbles, after which the mouse was returned to home-cage and number of marbles buried (min 2/3) was counted. Experimenter was blinded to treatment groups. For the CFA mice, the experiment was run in two different cohorts, by two different experimenters, with 10 months between, and showed the same pattern.

**Elevated plus maze.**
Animals were habituated to the experimental room for a minimum of 60 min before initiation of the experiment. Depending on group, mice were injected with saline, PBS, or 10 µmol/kg mPD5 (10 µL/g) and left in their home-cage. One hour later, mice were placed in the center of the elevated plus maze facing a closed arm and left for five min. Maze was rinsed with water between mice, and video recordings of the five min were analysed by EthoVision. Experimenter was blinded to treatment groups. For the CFA mice, the experiment was run in two different cohorts, by two different experimenters, with 10 months between, and showed the same pattern.

**Spatial learning and memory.**
On each day, animals were moved from the housing facility to the experimental room and allowed to acclimatize for at least 60 min before initiation of the experiment. All drugs used (PBS or 10 µmol/kg mPD5) were injected s.c. in a 10 µl/g body weight volume 60 min before first placement on the Barnes' maze on day 1-9 of the experiment. The experiment was divided

into Barnes maze habituation, spatial acquisition (training) and probe trials. During *habituation to the Barnes maze* itself, animals were placed under a cylinder in the middle of the Barnes maze for 10 seconds and then gently guided to the escape box and left there for two min. During *spatial acquisition (training)*, each mouse was placed under a cylinder in the middle of the Barnes maze for 10 sec followed by up to three min of free exploration of the Barnes maze. Once the animal entered the escape cage, it was left there for one min before being transferred back to its home cage for 15-20 min. This training was performed four times per day per mouse. During *probe trials* the reference memory of the animal was tested by placing each mouse under a cylinder in the middle of the Barnes maze for 10 sec followed by 90 sec of free exploration of the Barnes maze with no escape cage. On days 6-8, the escape hole was moved 90 degrees, and all mice went through the second bout of training. On day nine, a second probe trial was performed testing the reversal learning of the mice.

**Assessment of limb use.**

Cancer induced bone pain was induced as described in the "**Induction of cancer induced bone pain (CIBP model)**"-section. For the assessment of limb use, all mice from one cage were placed in an empty transparent plastic box and left for ten min to habituate. One mouse at a time was moved by tunnel handling or scooping to an identical plastic box for the limb use test. The gait of the mouse was observed for three min and scored by the following system: 4 Normal gait, 3 Insignificant limping, 2 Significant limping and shift in bodyweight towards the healthy limb 1 Significant limping and partial lack of use of ipsilateral leg 0 Total lack of use of ipsilateral leg. A score of 0 was used as a humane endpoint. Urine and faeces were cleaned from the plastic box with a dry paper towel between mice from the same cage, and a new plastic box was used for mice from different cages. The limb use test was video filmed for later evaluation if necessary.

**Assessment of static weight bearing.**

Cancer induced bone pain was induced as described in the "**Induction of cancer induced bone pain (CIBP model)**"-section and the static weight bearing test was performed using an incapacitance tester (Incapacitance Tester, Version 5.2, Linton). Animals went through 3 equipment habituation sessions before baseline testing. For the weight bearing measurements, the mouse was guided into the tube in the correct diagonal position forcing its hind paws to be located on the two individual sensor plates. The sensor plates measured the applied force by the left and right hind legs respectively. The force was measured for three seconds where the mouse should remain still in the same position. While taking the measurement the tail was lifted gently from the sensor plates to avoid interference. Measurements were done in triplicate, where the mouse was removed and reintroduced into the tube between measurements.

**Single exposure place preference.**

Initial perception of the drugs was measured by single exposure place preference experiments performed in an elongated three compartment apparatus (67.5 cm x 24 cm) with a biased design of different floor textures and wall patterns and a neutral zone in the middle. We have previously shown that mice of both genders exhibit a strong bias towards the striped compartment and that a single exposure of psychostimulants in the grey compartment sufficiently changes the preference towards that compartment (9-11). For further details of the setup, we refer to (10). Experiments lasted three days with exposure sessions on days one and two and a preference test on day three. On each day, animals were moved from the housing facility to the experimental room and allowed to acclimatize for at least 60 min before initiation of the experiment. mPD5 was paired with the grey compartment, known to be the least preferred compartment (10, 11). All mice from a cage were tested at the same time, but not all were given the same treatment. Mice were weighed and injected s.c. with PBS or 30 μmol/kg mPD5. For the sePP on CFA-injected mice, the injury was induced as described in "**Induction of inflammatory pain (CFA model)**"-section. sePP was run on day 3-5 after CFA-injection, and von Frey was performed on days 0, 2, and 5 as described in the "**Mechanically-evoked pain (von Frey)**"-section.

**Conditioned place preference.**
CPP was performed in Med-Associates square behavioral chambers fitted with beam-break movement detection. Chambers are split in two (27.3 x 13.5 cm each) with a red Plexiglas wall (allowing penetrance of the beams) and a predictable-preference design. Least-preferred compartment: white walls + floor and a transparent lid. Preferred compartment: horizontal grey and black stripes on the wall and lid with a light grey upside-down Lego® floor. Mice can freely move between the two compartments through a 4x4 cm opening in the separating wall during pretest and posttest but are confined to one compartment during conditionings.
*Pretest (DAY 1):* To test the basal preference of the mice, they were placed in the chamber to freely move between the two compartments for 30 min. Time spent in each compartment was recorded and showed a preference for the striped compartment in all mice. Time spent in each compartment was used to define three even groups.
*Conditioning (DAY 2-9):* Mice were injected s.c. with PBS, morphine (10 mg/kg) (RH Pharmacy, Denmark, Vnr. 475249), or 10 μmol/kg mPD5 immediately before being placed in and confined to the appropriate compartment for 40 min. Drug was paired to the least-preferred compartment, conditioning was alternated so that mice were never placed in the same compartment two days in a row, and the day on which they were exposed to drug was counterbalanced.
*Preference test (DAY 10)*: Mice were allowed to freely move between the two compartments for 30 min.

**Ultrasonic vocalisation (USV) recordings.**
Mice were left in Med-Associates square behavioral chambers (27 cm x 27 cm) to freely roam around for 60 min, four days in a row. The first two days were for habituation, but on the third and fourth days, USV recordings were performed using an Avisoft UltraSoundGate 416Hb recording interface (Model CM16-CMPA, Avisoft Bioacoustics, Germany) at a sampling rate of 250,000 Hz in 16-bit format. On day three, both control and SNI mice were placed directly in the chambers and recorded for 60 min. On day four, control mice were injected s.c. with PBS, and SNI mice were injected with either 10 μmol/kg mPD5 or PBS immediately before being placed in the chambers and recorded for 60 min. The recordings were analyzed using the Avisoft SASLab Pro software (Version 5.02.07, Avisoft Bioacoustics) as described in (12), applying a fast fourier transformation (FFT) (1024 FTT length, 100% frame size, Blackman window, and 87.5% time window overlap). Spectrograms were produced at a frequency resolution of 244 Hz and a time resolution of 0.512 ms. Spectrogram analysis was performed using an automatic whistle-tracking algorithm with an element separation threshold of -45 dB, 1 ms minimal duration, and a hold time of 10 ms. To compare pain-relevant vocalizations, only *short* and *constant* USVs of 37 kHz were counted (12, 13).

**Tail-Flick immersion test**
Naïve wildtype male and female mice (10 weeks of age) were habituated to handling and experimental set up for one week prior to data collection. On the day of data collection, a baseline measurement was performed prior to injection of compounds. One measurement consisted of immersion of 2/3$^{rds}$ of the tail in water (49 °C) for maximum ten seconds, three times with an interval of ten seconds between each measurement using the mean of the three

measurements. After baseline, mice were injected s.c. with mPD5 10 µmol/kg mPD5, 10 mg/kg morphine (RH Pharmacy, Denmark, Vnr. 475249), or sterile PBS one hour before undergoing the tail-flick hot water immersion assay again.

**Capsaicin test**

Mice were pretreated with 10 µmol/kg mPD5, 10 mg/kg morphine (RH Pharmacy, Denmark, Vnr. 475249), or PBS 45 min before habituation to observation container (glass cylinder surrounded by mirrors on 2 sides) for 15 min. Mice then received an intraplantar injection of 20 µl capsaicin (1.6 ug/paw) (sigma CAS Number:404-86-4, diluted with 1:1:8 solution of tween-20, 70% ethanol, and MilliQ water) in one hind paw and were reintroduced to the observation container. Mice were recorded for three min after i.pl. injection with capsaicin and euthanized the same day. Mice were scored manually for the amount of time spent paw licking within three min after capsaicin administration.

**Fertility test**

Mice were 10 weeks old at the beginning of the experiment and acclimatized to the animal facility for two weeks after arrival. Male mice were randomized into two groups (n = 8/group) and weighed and injected s.c. once daily (between 3 pm and 4 pm) for 14 days with sterile PBS or 10 µmol/kg mPD5 (diluted in sterile PBS and pH adjusted to 7.4). After 14 days of dosing, the males were paired with female mice in fresh cages, and the number of pubs counted for 35 days (males were removed from the breeding cages after 14 days). Following the 14 days mating period, males were sacrificed by cervical dislocation caudae epidymidis were removed and weighed. Caudae epidymidis were cut into smaller pieces in 2 mL PBS (37 ºC) and incubated (37 °C, 5% $CO_2$) for 15 min. Following incubation, the mix was swirled gently, and 0.5 mL of the liquid was transferred to room tempered PBS to achieve a final volume of 1 mL. Subsequently, sperm count was determined by transferring 50 µL of the sample to a hemocytometer, and each sample was determined in duplicate by an experimenter blinded to drug treatment. Following criteria were assessed: mobility (forward and circular), local tail mobility and total number. Females were weighed on a regular basis from removal of the males until after birth.

**Statistics**

Unless otherwise stated, all data were analysed using GraphPad Prism 9.4.1. and presented as mean +-SEM with the significance level set to $p < 0.05$. For locomotion and elevated plus maze, Ethovision XT 13 (Noldus) was used to track the distance travelled by each animal. For within group analysis (i.e., von Frey experiments), two-way ANOVA followed by Dunnett's post hoc test was used. For between group analysis of multiple time points (i.e., locomotion experiments), two-way ANOVA was used without post hoc test due to lack of treatment significance in the ANOVA test. For the limb use experiment, data pre versus post treatment was analysed with a Kruskal Wallis test due to the non-parametric nature of the data. When analysing histograms of three groups or more (i.e., marble burying and elevated plus maze), one-way ANOVA followed by Tukey post hoc test was used. When analysing histograms of two groups (i.e. single exposure place preference), unpaired t-test was performed. ROUTS outlier analysis was used for the elevated plus maze identifying 2 outliers in the CFA dataset, that were removed from the data analysis. The experimenter was blinded to treatment throughout all experiments. Results on statistics can be seen in Supplemental Table S3.
